# Testing Finch’s hypothesis: the role of organismal modularity on the escape from actuarial senescence

**DOI:** 10.1101/771378

**Authors:** Connor Bernard, Aldo Compagnoni, Roberto Salguero-Gómez

## Abstract

1. Until recently, senescence was assumed to be a universal phenomenon. Evolutionary theories of senescence predict that no organism may escape the physiological decline that results in an increase in mortality risk and/or decline in fertility with age. However, evidence both in animals and plants has emerged in the last decade defying such predictions. Researchers are currently seeking mechanistic explanations for the observed variation in ageing trajectories.
2. We argue that the historical view on the inevitability of senescence is due, in part, to the development of its classical theories, which targeted primarily unitary organisms. In unitary species, the integration of resources and functions is high, and adult size is determined. In contrast, the architecture of modular organisms is indeterminate and built upon repeated modules. The isolation of mortality risk in species like hydra (*Hydra spp.*) or creosote brush (*Larrea tridentata*) may explain their null or even negative senescence.
3. Caleb Finch hypothesised three decades ago that species with the ability to compartmentalise risk may escape senescence. Here, we first review the evidence on organisms that slow down or even avoid senescence in the context of their architecture, along a continuum of unitarity-modularity. Then, we use open-access databases to comparatively analyse various moments of senescence and link longevity to the degree of anatomic modularity. Our analysis compares 138 plants and 151 animals. Our comparative analysis reveals that plant species that are more modular do indeed tend to escape from senescence more often than those that are unitary. The role of modularity in animal senescence is less clear.
4. In light of novel support for Finch’s hypothesis across a large diversity of plant species, and with less conclusive findings in animals, we identify new research directions. We highlight opportunities related to age-dependent mortality factors. Other areas for further research include the role of modularity in relation to endocrine actions, and the costs of modular anatomies.

> *“The actinozooid is a living thing which knows no time of youthful vigour, no waxing to a period of adult life, no waxing to senility – it knows no age – it practically knows no natural death.”* – Wood-Jones (1912)

## 1 THE EVITABILITY OF SENESCENCE

Since the late 1970s, three major theories have persisted to explain declines in physiological performance with age, that is *senescence*. These are: (1) Medawar’s (1952) theory of mutation accumulation, whereby organisms senesce due to their inability to revert the rate of accumulation of deleterious mutations; (2) Williams’ (1957) theory of antagonistic pleiotropy, whereby genes with dual effects (with positive effects of fitness early in life but negative late in life) are selected because “the young matter more than the old”; and (3) Kirkwood’s (1977) disposable soma theory, whereby organisms with a clear physical separation between germ and soma lines senesce because natural selection selects the germ over the maintenance of the soma. Medawar’s and Williams’ theories center on genetic principles for a discounting of late-life performance (Le Bourg 2001; Promislow et al. 1999; Hamilton 1966). Williams’ and Kirkwood’s theories share ground in staking their respective claims on ties between fundamental biological constituents. These two theories identify links between between genes’ actions (*i.e.,* early *vs.* late) and between functional allocations (*i.e.*, maintenance *vs*. reproduction), respectively (Turke 2008; Kirkwood and Austad 2000; Kirkwood and Rose 1991; Rose 1991). The three classical theories of senescence aim to identify the forces responsible for shaping patterns of senescence. However, they largely fail to explain how or under what conditions senescence could be postponed or even reverted.

In the decades since the classical theories for senescence were posited, research has qualified the apparent inescapability of senescence. For example, Hamilton (1966) asserted that actuarial senescence necessarily follows from the exponential growth of population. He mathematically showed that organisms with demographies highly favorable to late-life performance still experience selection bias for early-life performance. Implied here is that populations with non-exponential growth could escape senescence. Williams (1957) described senescence as a transient genetic phenomenon in which genes that present mixed benefits are partially substituted by more adaptive genotypes. According to this theory, variation in the genes and in their influence on one another can generate positive, negligible, and negative senescence. Negative senescence describes a decline in mortality after maturity (*sensu* Vaupel 2004).

More recent models of senescence describe how density dependence can accelerate or decelerate senescence as a function of stage-dependent mortality (Abrams 1993). Other researchers (Shefferson, Jones, and Salguero-Gómez 2017; Gómez 2010; Turke 2008; Buss 1987) have noted another qualification, implied by Williams’ work based on cell differentiation. Organisms with undifferentiated cells (thus without soma-germ separation) may not be subject to patterns of senescence assumed under all classical theories for senescence. Surprisingly, the existence of alternative explanations for escaping senescence did not detract from Hamilton’s (1966) prediction of universal senescence until the most recent literature (Nussey et al. 2013; Baudisch 2005).

A systematic review or estimate for how many taxa are accountable to the above-mentioned exceptions does not exist, that we are aware of. We also do not know how the closely proposed mechanisms that release species from senescence relate to particular groups of species. This information can advance our understanding of how much has been explained under existing theory, and can focus future studies. While the conceptual foundations of these exceptions are strong, they have rarely been tested empirically. For example, Vaupel et al. (2004) offer a cogent proof of negative senescence based on growth-related functional performance. They ground theoretical advancements with examples of species that exhibit negligible or negative senescence that are also indeterminate in their growth. Until recently, we were not aware of any work connecting how closely indeterminate growth predicts negligible senescence, nor analysing the existence of confounding influences. Early studies are now underway (Cousin & Salguero-Gómez, *pers. comm.*).

Taxonomic bias in early studies of senescence in unitary organisms (e.g. mammals, birds) likely contributed to the notion that senescence is universal (Jones et al. 2014; Monaghan et al. 2008; Shefferson et al. 2017). The idea of universal senescence may also be attractive because cellular ageing, not far afield of senescence, is written about in universal terms (*i.e.*, cellular senescence; noted by Comfort 1979). Statements in recent publications put little distance between cell and organismal finitude: “Organisms wear out, just like machines” (Ricklefs 2008); “[D]amage will accumulate in parallel with cells” (Kirkwood 2005); “Aging and its associated decline in survival and reproductive fitness is an inherent feature of biological systems” (Alper, Bronikowski, and Harper 2015). Studies on ageing focus on a set of cellular and molecular processes that are distinct from those that are the focus of senescence. Ageing processes include telomere shortening (Young 2018); oxidative stress (Monaghan, Metcalfe, and Torres 2009); double-strand break (White and Vijg 2016); and related mechanisms underlying the damage or error theories of aging (Jin 2010; reviewed in connection with senescence by Comfort 1979; Petralia, Mattson, and Yao 2014). Insofar as variations in the ageing processes at the molecular level influence the types and rates of age-dependent physiological degradation, they may be an important dimension to the discourse on senescence.

The extent to which ageing processes should be treated independently of senescence, evolutionarily and mechanistically, is contested. The distinction between the two relates to a distinction in evolutionary theory related to evolutionary constraints (Wrycza, Missov, and Baudisch 2015; Wensink, Caswell, and Baudisch 2017). The approach we use, based on Baudisch (2011), negotiates this conflict by separating two central axes of variation –between pace and shape. This approach enables a clean differentiation between patterns of life expectancy and mortality patterns.

We argue that the inevitability of organismal senescence should hold only so far as molecular declines upscale to the demography of the organism. This upscaling –from cellular degradations to physiological injury– must go through the anatomy (*i.e.,* tissues) and physiology (*i.e.* organs) of the individual. Considering this up-scaling of cellular declines to physiological declines may inform the mechanisms for relieving senescent forces of natural selection.

Recent research has highlighted an important aspect that was absent from whole-organismal senescence with respect to plants and non-unitary animals (Jones et al. 2014; Vaupel et al. 2004; Gardner and Mangel 1997). Many cases of negligible senescence (*i.e.*, mortality [Fig. 1b & e] and/or fertility [the latter not addressed analytically in this review] remaining invariable with age) or even negative senescence (mortality risk decline [Fig. 1e & f] and/or fertility increase with age [again, the latter not addressed in this review, but see Baudisch and Stott 2019] are now known based on reliable data (Pletcher, Houle, and Curtsinger 1998; Finch 2009). Perhaps not by coincidence, these patterns occur in organisms not studied (or even discovered!) when the classical theories of the evolution of senescence were developed. Some of these include Brandt’s bat (*Myotis brandtii*; Podlutsky et al. 2005), barn owl (*Tyto alba*; Altwegg, Schaub, and Roulin 2007), greenland shark (*Somniosus microcephalus*; Nielson et al. 2016), rockfishes (Cailliet et al. 2001; Mangel, Kindsvater and Bonsall 2007; Munk 2001), ants and termites (Carey 2001), naked mole-rat (*Heterocephalus glaber*; Buffenstein 2008), *Hydra* (Schaible et al. 2015; Martínez 1998), bristlecone pine (*Pinus longaeva*; Lanner and Connor 2001), *Borderea pyrenaica* (García, Espadaler, Olesen 2012), and wilcox brush (*Eremophila forrestii*; Erlén and Lehtilä 2002).

**Figure 1.**
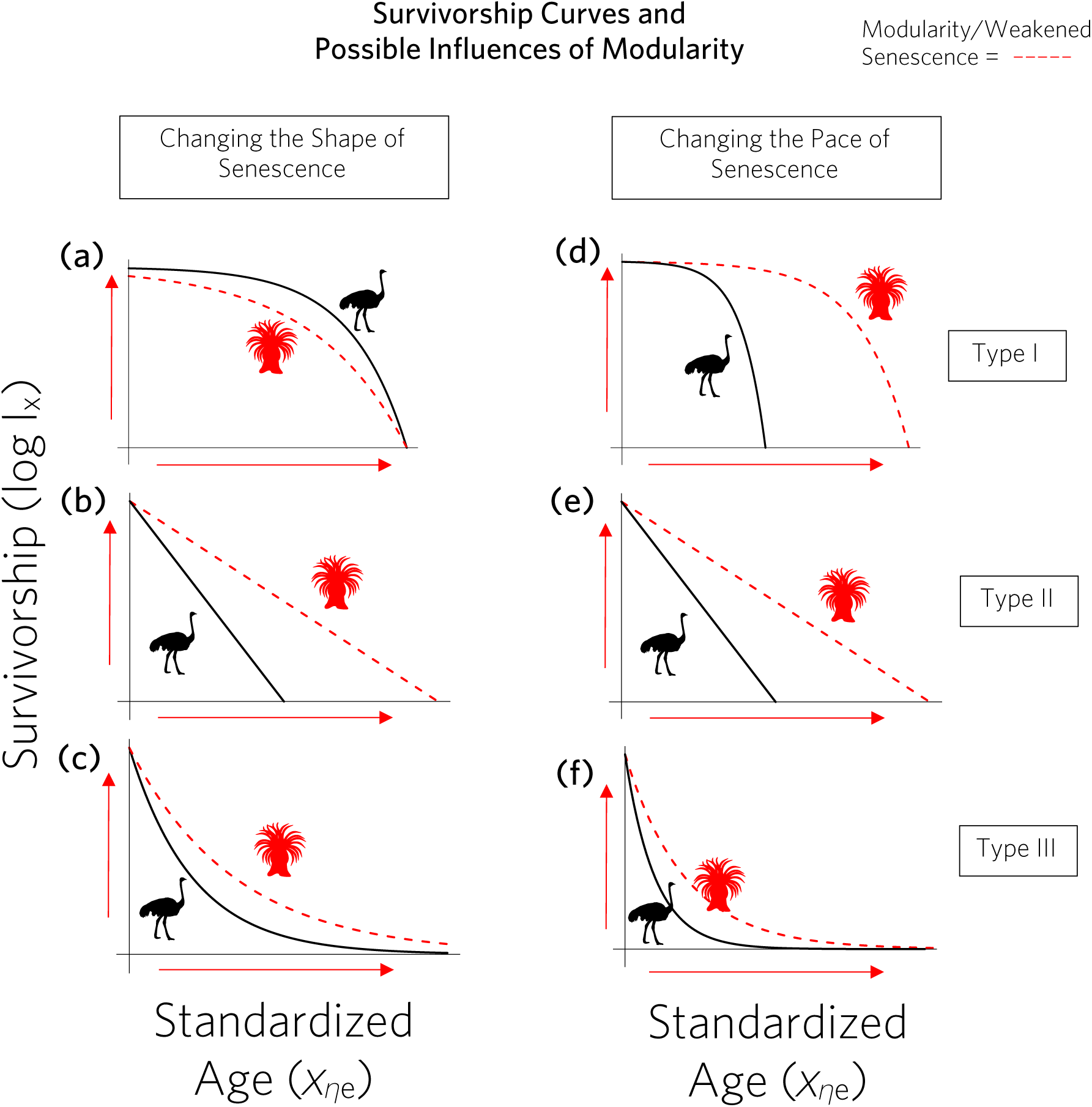
A graphic representation of Finch’s Hypothesis for actuarial (*i.e.,* mortality) senescence. These figures represent the hypothesised trait-senescence relationship in which modularity will lower the shape (a, b, c) and reduce the pace (d, e, f) of senescence, extending life span and reducing late-life mortality. The black lines labeled with ostrich icons represent a hypothetical unitary organism (with high physiological integration), and red lines labeled with coral icons represent a hypothetical modularised organism (with low physiological integration). Finch’s hypothesis predicts that more modularised physiologies (the red lines) should result in reduced senescence (*i.e.*, longer lifespan and lower late-age/stage mortality) relative to their unitary counterparts (black lines). The survivorship curves are characterised by survivorship type (Type I: a, d; Type II: b, e; and Type III: c, f), reflecting standard groupings of mortality patterns, which is discussed further in Figure 2. Note that changes in the shape of a Type II curve obligates either a Type I (a, d) or III (c, f) curve, but pace can drive a less severe slope, which takes its place in this graphic. Also, note that the difference in shape and pace curves among Type 3 (c, f) curves are more subtle than for Type I (a, d) curves; the difference between panels are most noticible in how they intercept the x-axis. If the effect of modularity on senescence were strong enough, a species or lineage could be predicted to pass from one survivorship type to another – from Type I, to II, to III, respectively.

Caleb E. Finch, in his seminal monograph of senescence (1990), brought attention to the few known species that at the time suggested negligible or even negative senescence. In this monograph, Finch posited a novel connection between senescence and anatomy, by probing the question of how modular forms could alter senescence. Finch suggested that if modules age independently or incur injury independently, as physiological units, then they may insulate the composite individual from those effects. Thus, Finch inquired whether modules could lengthen the life of an individual by preventing local injury from translating to injury of the individual. Until recently, testing this hypothesis has been hindered by a dearth of relevant data and substantial literature across many species. Here, we avail of recent advances in histology and physiology to proxy the degree of modularity, and evaluate links between demographic data and actuarial senescence.

Here, and in line with classical use, we use the term “modular” to describe individual organisms that are constructed of multiple, repeat sub-units that are individually dispensable without injury to survivorship of the individual (Vuorisalo and Tuomi 1986; Vuorisalo and Mutikainen 1999). Within this framework, the individual can be thought of as a grouping of genetically identical cells which share physiological interdependence (extending from criteria distinctions discussed by Pepper and Herron 2008; Pradeu 2016; DiFrisco 2017). This usage allows for minor genetic variation that occurs from random, somatic point mutations. Clones are therefore considered parts of a single individual –as are modules– despite their heightened degree of physiological independence and compartmentation. Alternative ways to define individals may be used on the basis of fundamental evolutionary units, specific physiological properties, or other factors. However, our definition facilitates comparison across broad morphological types and taxonomies, while retaining a demographically coherent unit. Our analysis shows that traits that reflect a higher degree of physiological modularity correlate with negative senescence in plants, although not necessarily in animals. These findings therefore provides novel empirical support for a possible explanation for escaping senescence.

## 2 A COMPARATIVE OVERVIEW OF FINCH’S HYPOTHESIS

We extend Finch’s hypothesis by linking its predictions to the *pace* and the *shape* of senescence. The distinction between the pace and shape of senescence emerged long ago (Keyfitz 1977) as a useful framework in ageing research that decouples important metrics of ageing (Keyfitz 1977; Baudisch 2011; Baudisch et al. 2013, Wrycza and Baudisch 2014). The pace of senescence represents the characteristic mean life expectancy (*η_e_*) of an organism. For example, the lifespan of moss campion *(Silene acaulus*; routinely >50 yrs, some specimens >300 yrs; Morris and Doak 1998) is greater than that of the invasive cheatgrass (*Bromus tectorum*; <1 yr; Compagnoni and Adler 2014). On the other hand, the shape of senescence describes *when* and how intensively mortality hazards change during the *normalised* lifetime (typically over *η_e_*; Keyfitz 1977) of a species (Fig. 2). Because shape is normalised (*i.e.,* age-value divided by mean life expectancy), it allows one to compare the rate of senescence across many species, regardless of differences in *η_e_* across populations and species.

**Figure 2.**
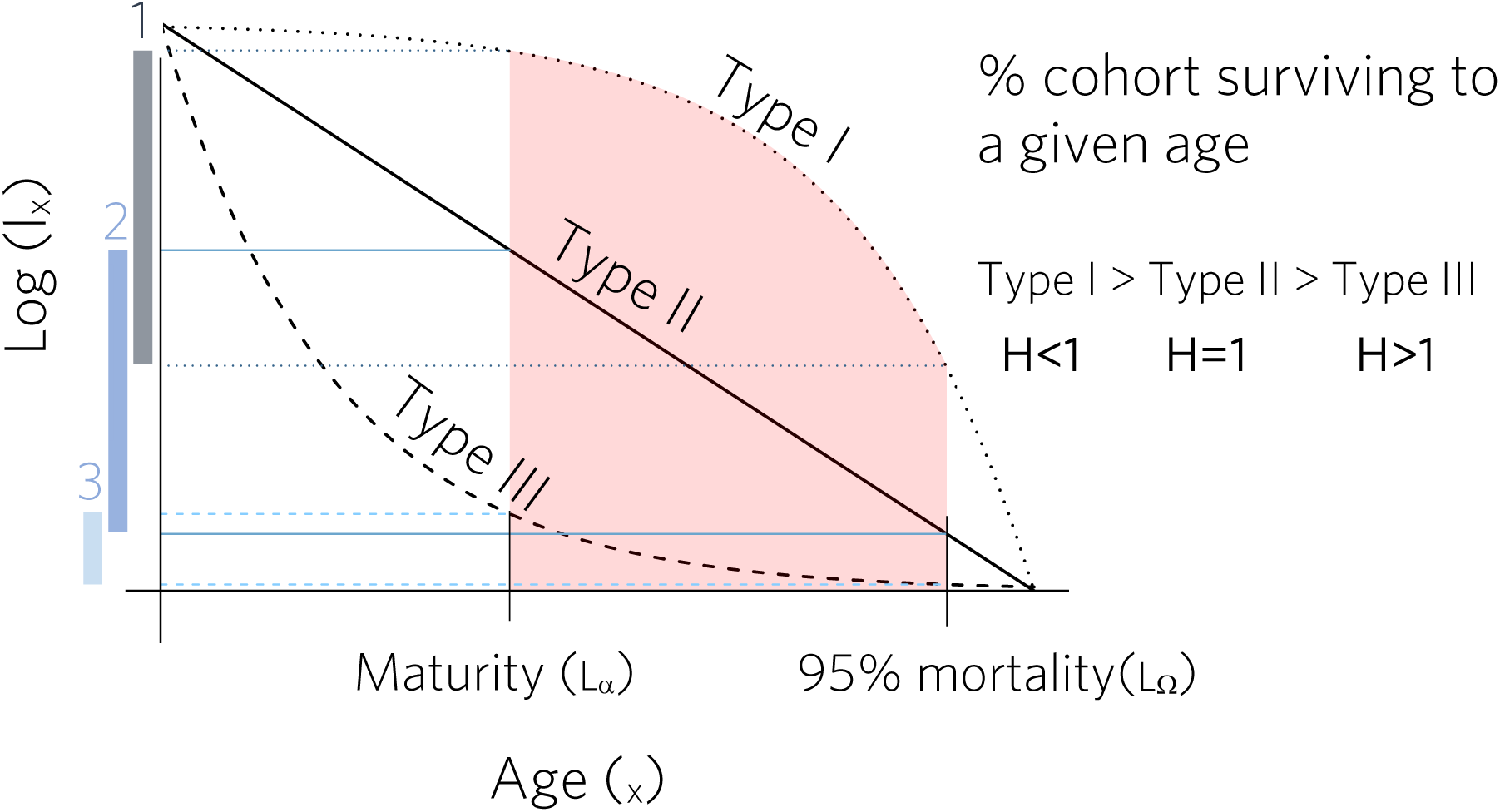
A graphic representation of Keyfitz’ entropy (*H*). Rates of actuarial senescence can be measured across a wide range of dissimilar actuarial dynamics (in our case, examined across 138 plant and 151 animal species), using a standardised metric of *H*. Keyfitz’ entropy *H* quantifies when mortality occurs along the lifetime of a cohort, from age at maturity (L_α_), until –in this case– 95% of a cohort has died. This metric is dimensionless and normalised by the mean life expectancy of the organism, which allows for intra- and inter-specific comparisons, and is directly linked to the degree of modularity of different organisms (See Figure 3). Survivorship types describe different mortality patterns with age. In Figure 3, which highlights a period from maturity to 95% cohort mortality in red, a larger fraction of total mortality occurs in Type I and type II mortality than in Type III, as shown by the length of bars on the y-axis, which correspond by color to the survivorship lines. To a lesser extent, more mortality occurs in type I than type II over this same interval. The normalised metric represented by bar length (percent cohort change) indicates the mortality change over the *entire* interval; however, this measure of change can be measured at any age interval within this range. *H* is a metric of whether mortality through time tends to be weighted in early life, nearer to maturity, or late in life, nearer to 95% cohort mortality. Accordingly, Type I survivorship has the greatest *H*, with higher mortality change occurring late in the interval, near *L_Ω_*, and Type III has the lowest *H*, with mortality change occurring late in life, near *L_α_*.

Following Finch’s hypothesis, we expect greater functional modularity to negatively influence the pace, and, to? a more limited degree, the shape of senescence. Modularity should affect the pace of senescence through the individual’s ability to minimise damage by shedding damaged modules (Kozlowski 1973; Denley and Metaxas 2016). Examples of modules that could be shed include specific ramets (in clonal organisms), wood vessels, cnidarian polyps, bryozoan polymorphs, and ascidian zooids. In general, we do not expect the shape of senescence of an individual to be influenced by modularity. This is because neither theory nor evidence suggest that localised injury is linked to age-specific mortality any more or any less than other mortality factors. However, modularity might alter shape if the accrual of mutations (*sensu* Medawar 1952) occurs on a modular basis and age-dependent declines are restricted to the module. Because multiple prerequisites underlie this proposition, we consider it a subsidiary hypothesis.

The scope of our review encompasses multiple kingdoms and a wide array of taxonomic groups. By approaching the commonalities of modularity broadly, the structural appearance and functional role of modules is diverse across different species. For example, plants can have deeply modularised anatomies where physiological independence among structurally discrete components of an individual is high (e.g., ramets of an aspen grove). In contrast, animals tend to be “unitary” individuals, meaning that they have structural anatomies where physiological integration is high across their body plan. In these organisms, “shedding” part of a unitary animal (e.g., loss of an appendage or loss of a segment of the body) tends to be, by definition, physiologically disruptive to the individual.

The potential for greatly different physical and functional qualities of modules is unified by a basic characterisation: modules need not be inclusive of the full range of physiological functions carried out by the individual such as water balance or energy regulation. Instead, modules can be recognised as repeat sub-units within just one of those physiological functions (e.g., immune activity, renal activity, respiratory activity). This latter point suggests that we can divide our analysis into kingdoms, separating it between unitary (animals) and non-unitary (some animals like corals and sponges, and plants) organisms.

In general, traits that enhance modularity should correlate positively with the pace of senescence (Silvertown, Franco, and Perez-Ishiwara 2001). In plants, we expect increases in clonality, stem sectoriality (Schenk et al. 2008; Ewers et al. 2007), and bark stripping (*i.e.*, the ability of trees to persevere when only a small part of the trunk is surrounded by live cambium, Matthes et al. 2002) to prolong longevity. In animals, we expect the number of polyps within a colony (Denley and Metaxas 2016) and the number of functionally redundant organs to prolong longevity. The number of discrete, redundant pathways for blood or lymph circulation (Seiler 2010; Huang, Lavine, and Randolph 2017 respectively), and the number of parallel endocrine pathways should do the same. We expect that all of these traits traits should correlate negatively with mortality rates, and therefore increase the pace of senescence (Fig. 1a, b & c).

On the other hand, the link between modularity traits and the rate of mortality over age (*i.e.*, the shape of senescence) is subtler. In plants, several traits could change age-specific mortality rates, including clonality, epicormic branching in trees, and indeterminate growth. On the other hand, in animals, fewer traits would be expected to change age-specific mortality rates, because growth tends to be definite. A notable exception to this is metazoans (Bodnar 2009). The indirect effect of age-related size-dependent benefits can potentially tap alternative mechanisms for escaping senescence detailed by Vaupel et al. (2004).

Modularity is expected to decrease the pace of senescence, with possible indirect effects on the shape of senescence. As discussed above, modularity may indirectly affect the shape of senescence as a result of facilitating the attainment of advanced ages, and size-dependent benefits. More generally, modularity could have a more complex effect on the shape of senescence if localised injuries affect age-specific mortality. Some mortality factors are internally mediated and dependent on lifespan-normalized age; others are external and independent of age. As a result, if modularity facilitates the attainment of more advanced ages, we would expect that the pertinent mortality factors will change. These changes may alter when in the lifespan of an individual mortality is most likely to occur, and thus change the shape of senescence.

This relationship is captured in the generalised age-dependent/degenerative term under the Reliability Theory of Senescence (Gavrilov and Gavrilova 2001). Where environmental mortality factors influence mortality rate irrespective of physiological declines, they are mapped as age-independent mortality factors. Under Reliability Theory, modularity will influence the realisation of a system failure (which we define at the scale of the individual), as it constitutes a system component with functional redundancy that, in turn, acts as a redundancy parameter, reducing *failure events:system failure* (*i.e.*, local injury:individual mortality).

### 2.1 Modularity and senescence in plants

Several lines of evidence suggest that structural characteristics of plants that increase their modularity also lengthen their longevity. One line of evidence relates to epicormic branching in trees – the ability to produce branches directly from sections of the trunk (Meier, Saunders, and Michler 2012). Epicormic branching has been linked to longevity in long-lived trees, such as the Douglas fir (*Pseudotsuga menziesii*; Ishii et al. 2007). Epicormic branches of the Douglas fir allow old, large trees to maintain foliage (Ishii et al. 2001), which is crucial to provide the large photosynthetic capacity that allows the trees to maintain their characteristic high growth rates at advanced ages (Sillett et al. 2010). Moreover, modularity clearly promotes large sizes in many clonal species of plants. Some of the largest and oldest individual organisms on earth are clonal plants, such as the Pando aspen clone (*Populus tremuloides*; DeWoody et al. 2009; Mock et al. 2008) or a Neptune grass clone (*Posidonia oceanica*) found in the Mediterranean Sea (Guerrero-Meseguer, Sanz-Lázaro and Marín 2018).

Demographic effects of physiological injuries also appear to be informed, at least in part, by the structural organisation of an organism. Severe, long-term impairments of a physiological system can be fatal for some species and merely locally-injurious, and without individual-scale physiological injury, for others (Kozlowski 1973; Winston 2010). Whether one is injured can be a function of the degree of integration within the individual. The transport of water in plants is instructive in this regard. Since the 1980s, the formation of air pockets in hydraulic conduits within plant vascular tissues –*air embolism/cavitation*– has been found to be a prevalent cause of mortality in terrestrial plants (Tyree and Sperry 1989; Zimmerman 1983). Variation in the morphology that influences embolic risk is expected to have a high demographic elasticity.

The interaction of emboli with the morphology of plant vascular and vessel morphometry has been the focus of much research (Sperry et al. 2007; Venturas, Sperry, and Hacke 2017; Pitterman et al. 2010; Tyree and Sperry 1989; Ewers 1985; Bailey 1916). The well-established “rare pit hypothesis” of trade-offs between efficiency and safety of vessels stems from this line of questions (Christman, Sperry, and Adler 2009; Christman, Sperry, and Smith 2012). Of direct importance to senescence, the safety-efficiency vessel trade-off offers a case example of direct importance to senescence. When an embolism forms in a vessel, the vascular configuration of the plant determines whether and to what extent the initial embolic disruption spreads to the whole organism (*i.e.* runaway embolism *sensu* Sperry and Pockman 1993). More sectorial vasculatures can mitigate the risk of runaway embolism by reducing the lateral connections between vessel bundles. If emboli result in runaway embolisms, they would be contained to a few bundles (Zanne et al. 2014; Orians, Babst, and Zanne 2005).

A series of studies have explored the environmental correlates of this type of sectoriality, and have found higher sectoriality in habitats with higher embolic risk, notably deserts (Maherali, Pockman, and Jackson 2004; Wright et al. 2004; Ewers et al. 2007; Royer et al. 2005; Ehleringer 1980; Carlquist 1975; see, too, related questions addressed by Adler, Sperry, Pockman 1996). Indeed, Schenk and collaborators (2008) showed that sectoriality increases on either hemisphere towards the latitudinal belt of arid zones across 12 woody species.

To test the effect of modularity on senescence in plants, we quantified the correlations between measures of the pace and shape of senescence and proxies to plant modularity. We quantified pace and shape of senescence using the COMPADRE Plant Matrix Database (Salguero-Gómez et al. 2015) and used methods described in Salguero-Gómez (2016) and Jones et al. (2014). We implemented a method described in Baudisch et al. (2013) to compare inter-specifically various moments of senescence across species. This approach isolates measures of lifespan from lifespan-standardised metrics of survivorship, and reduces highly variable mortality patterns to select statistic features whose discrete markers reflect key variations in population structure.

We derived, from the resulting 138 plant species in COMPADRE, three metrics of the pace of senescence. These metrics consist of mean life expectancy (*η_e_*), maximum longevity (*η*_max_), and generation time (*T*) using methods developed by Caswell (2011). We used a single metric to characterise the shape of senescence: Keyfitz’ entropy (*H*) (Keyfitz 1977; Baudisch 2011; Wrycza, Missov, and Baudisch 2015), which describes the overall shape of the survivorship curve (Fig. 2), such that when H>1, mortality –and thus the rate of actuarial senescence– decreases with age. Thus, under this metric, the decay in survivorship is quantified irrespective of reproduction.

The metric *H*, as well as those that quantify the pace of life (e.g., *η_e_*, *η*_max_, *T*), rely on age-structured demographic information. The underlying demographic information is often captured in a stage-structured format (as matrix population models). To derive age-structure from stage-structured models, we used methods from Jones et al. 2014 (based on mathematical proofs by Caswell 2001). These age-from-stage extraction methods use numerical simulations of the stage-structured matrix to determine a quasi-stable-state of a population distribution. This, in turn, yields a decaying survivorship *l_x_* curve from which our desired summary metrics can be extracted in an age-structured form. When simulation models converged to a stable stationary population distribution, they were not included in our data. This method and convergence selection criterion were also used on the data for animals (below).

To test whether functional correlates of physiological modularity inform the pace and shape of senescence, we drew on several open source databases. We collected functional trait data related to the degree of modularity from the FRED (Iversen et al. 2017), BIEN (Enquist et al. 2009), and TRY databases (Kattge et al. 2011), as shown in Table 1. These global trait databases are curated repositories of plant traits collected from a wide range of field studies, laboratory experiments, and observations. Here, we consider only those quantified under unmanipulated conditions. When multiple functional trait values were available per species, we computed the mean of each species-specific trait value. This was a straightforward mean, which we performed after making sure that identical values of a trait (e.g. root diameter of 0.4996 mm) were not doubles. To do so, we examined several instances of double values in each one of the three databases, and contacted database curators for feedback on the most suspicious cases. We present only results of our analyses where >20 species were present in COMPADRE and in the functional trait databases (Table S2).

**Table 1.**
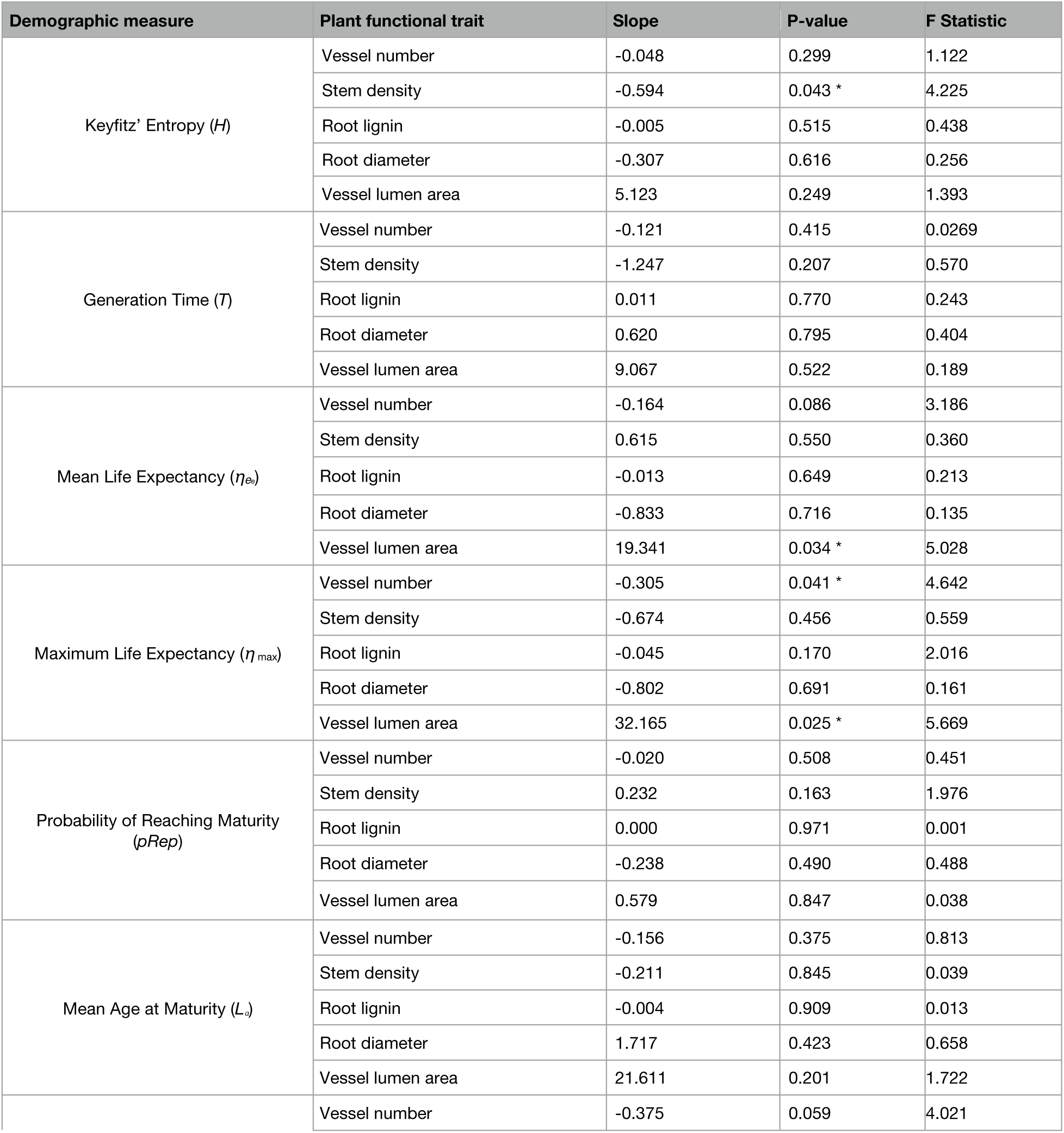
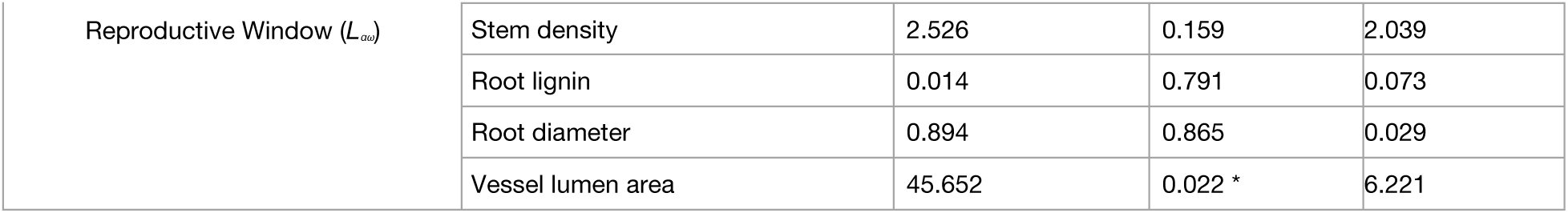
Full results of linear models of functional traits relating to metrics of senescence and longevity in plants. *: P<0.05.

**Table 2.** Full results of linear models of indices of anatomic modularity relating to metrics of senescence and longevity in animals. *: P<0.05; **: P<0.005; ***: P<0.001.

| Demographic measure | Animal functional trait | Slope | P-Value |
| --- | --- | --- | --- |
| Keyfitz' Entropy ($H$) | Renal Modularity Index | 0.046 | 0.019* |
| Keyfitz' Entropy ($H$) | Lymphatic Modularity Index | 0.001 | 0.967 |
| Mean Life Expectancy ($\eta_e$) | Renal Modularity Index | 1.636 | 0.274 |
| Mean Life Expectancy ($\eta_e$) | Lymphatic Modularity Index | 1.474 | <0.001*** |
| Maximum Life Expectancy ($\eta_{max}$) | Renal Modularity Index | 6.706 | 0.061 |
| Maximum Life Expectancy ($\eta_{max}$) | Lymphatic Modularity Index | -6.595 | 0.002** |

Finally, we tested for correlations between species-specific demographic traits and their modularity traits using a battery of phylogenetic generalised least squares (pgls) using the packages ape (Paradis, Claude, Strimmer 2004) and caper (Orme 2013) in R (R Core Team 2019). We used the phylogeny of vascular plants by Zanne et al. (2014). Family-wise errors were corrected using a Bonferroni correction. We used Aikaki Information Criterion (AIC) scores for model selection among a set of competing models with linear and quadratic predictors. The lowest AIC scores were linear for all models run and were therefore used to obtain our results.

### 2.2 Modularity and senescence in animals

The literature on senescence has historically been more inclusive of unitary animals than it has been for other groups. Important examples that have advanced our understanding of animal senescence include studies on roe deer (*Capreolus capreolus*; Gaillard et al. 1993, Loison et al. 1999); Soay sheep (*Ovis aries*; Catchpole, Morgan, and Coulson 2000); Dall sheep (*Ovis dalli*; Murie 1944; Deevey 1947; Kurtén 1953); yellow-bellied marmot (*Marmota marmota*; Berger et al. 2016); and the short-tailed vole (*Microtus agrestis*; Leslie and Ranson 1940; Leslie et al. 1955). However, to date, other lineages across the tree of life have not received much attention.

Research in senescence has only gained momentum in the Plantae Kingdom in the last two decades (Salguero-Gómez, Shefferson, and Hutchings 2013; Thomas 2013; Munné-Bosch 2008; Vaupel 2004; Thomas 2002) and it remains even less represented among modular animals (but see Bythell, Brown, and Kirkwood 2018 and Tanner 2001). Within plants, researchers have explored the question as to whether ramets senesce faster than genets (Salguero-Gómez 2017; Orive 1995). A similar angle on the animal kingdom remains untapped. This may be attributable to the close relationship of modularity to meristem-based growth, plant vasculature, and other functional characteristics restricted to the Plantae Kingdom (Vuorisalo and Mutikainen 1999; Clarke 2011). Broader application of these scaling relationships may also be hindered by the conflation of modularity with other coexistent functional units. Modules often coincide with carbon sectors, structural axes, independent physiological units, and inflorescences, among other such structures.

Here, we provide a review of animal modularity and its relationship to longevity and senescence. Our goal is to offer a translation of the primarily plant-based term modularity to individual animals for which the integration of resources is high and adult size tends to be determined. In doing so, we take a discrete step beyond previous treatments on the subject (notably Ewers et al. 2007; Vuorisalo and Mutikainen 1999; Harper 1980), in that we evaluate specific hierarchies of organ systems across demographic metrics.

At first sight, the term modularity would not be applicable to most animals if defined as repeat sub-units of multi-cellular tissue (Chapman 1981; Vuorisalo and Tuomi 1986; Finch 1990). Notable exceptions include corals and sponges, or, using a colony perspective, social animals such as ants (Kramer et al. 2015). Overall, however, the consideration of the Kingdom Animalia through the lens of classical modularity has to date received little attention. There is a corresponding lack of precision in describing the intertaxonomic concepts pertaining to modularity across kingdoms. The development of a framework that cuts across kingdoms allows us to test urgently needed mechanistic theories across the tree of life (Baudisch & Vaupel 2012).

A notable exception in this direction is the work by Esteve-Altava (2017), who recently interrogated a number of similarities in modular concepts. This research focused on structural, developmental, functional, and topological integration of modules in plants and animals, thus touching upon how we characterise modularity in this paper. However, in stressing the conceptual cross-over between plants and animals, Esteve-Altava highlights developmental origins, rather than the nexus of modularity with demography. The relationship between modularity and longevity found in plants (Fig. 3 a & b) raises the novel question of whether the degree of structural redundancy in animals is relevant to survival. Specifically, it raises the question of whether its effect on senescence is comparable to that of modules in plants. We tackle this question with a review of animal tissues and a criteria-based approach to qualifying modularity from a histological and functional angle.

**Figure 3.**
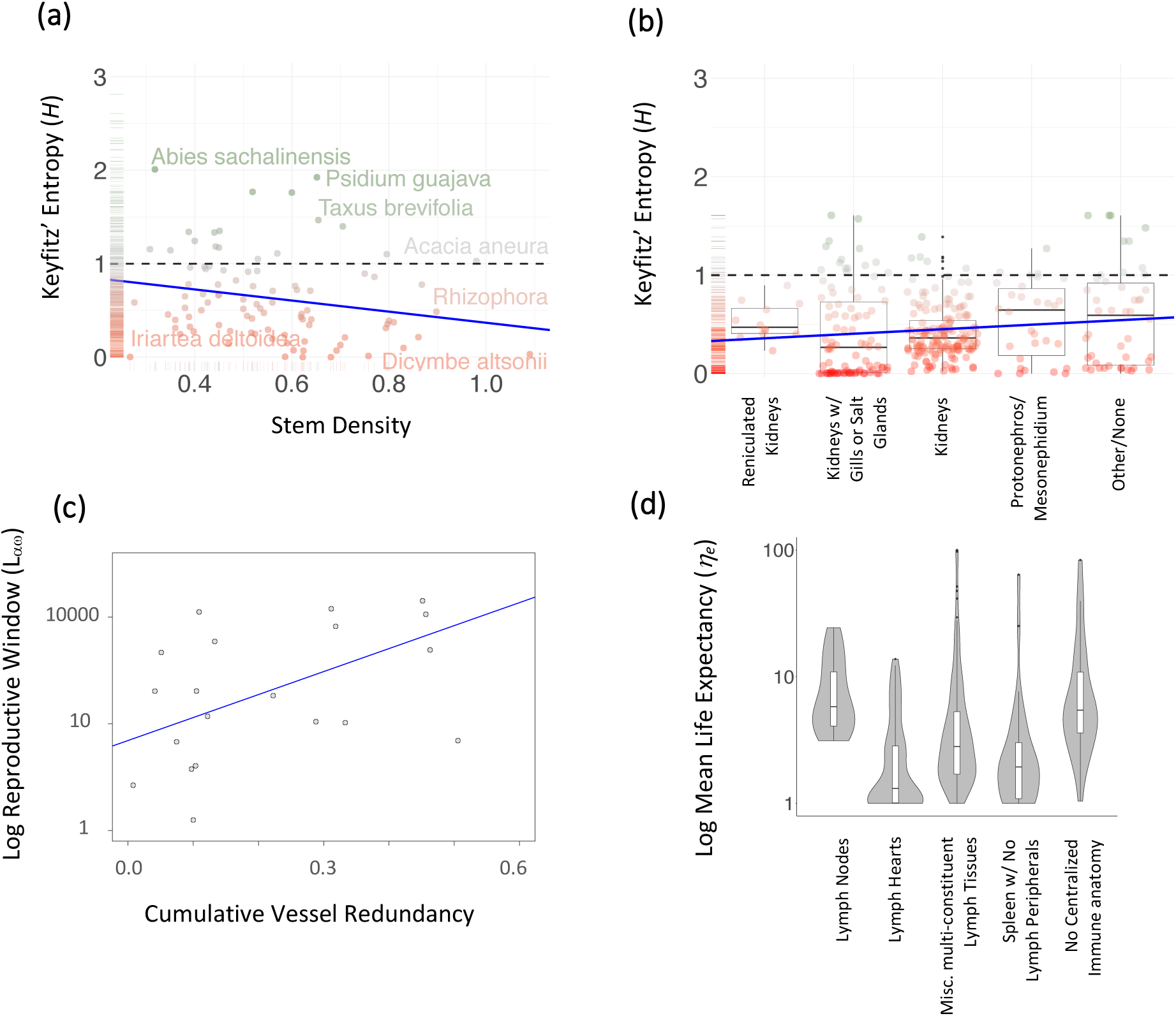
Finch’s hypothesis accurately predicts relationships between plant modularity as measured by functional traits and senescence, but fails to explain patterns in unitary animals between organ dividedness and senescence. Blue lines show significant relationships between the shape of senescence (Keyfitz’ entropy (*H*); see Figure 2 for more information on this metric) and (a) stem density, and (b) an index of renal modularisation. *H*=1, signified by the dashed black line, represents negligible or neutral senescence, and is the dividing line between positive and negative senescence. Larger *H*-values, higher on the y-axis (and color-weighted by green), represent lower rates of senescence; Smaller *H*-values, lower on the y-axis (and color-weighted by red), represent higher rates of senescence. While the stem-density effect on the shape of senescence supports Finch’s hypothesis, the renal index runs counter to expectation. (c) Cumulative vessel redundancy (lumen area corrected by vessel number) provides a measure of modular vessel units, quantified by number, corrected for vulnerability to embolism related to lumen area, and thereby is expected to be a more effective positive correlate of modularity than vessel number alone. Cumulative vessel redundancy positively correlates with reproductive window (*L_αω_*), consistent with the idea that modularised vessel anatomy slows the pace of senescence. (d) Mean life expectancy (*η_e_*) correlates negatively with an index of immune anatomy subdividedness, counter to the hypothesis that more sub-divided, and thereby more modularised, anatomy would promote longevity. There is wide variation in the distribution characteristics which map out differently among groups, though this is non-significant. Some of these characteristics include the lack of short-lived species with lymph nodes, and a tendency for higher representation of species with low lifespans in all groups except those with lymph nodes and no centralised immune anatomy. By reorienting the rank order of “no centralised immune system,” a stronger relationship would emerge.

The classical model of modularity (White 1979) describes modules along lines of discrete units of growth, units of reproductive independence (Harper and Bell 1979), independent functional units (Sprugel, Hinckley, and Schaap 1991), or independent physiological units (Watson 1986; Watson and Casper 1986). This principle works well to describe plants and colonial invertebrates, since these are comprised of well-defined, repeat structural sub-units. In other ways, it is deficient and unduly restrictive.

Modularity is a physiologically defined term whose touchstone is partial independence of different components of an individual. In most unitary animals, modularity does not appear to have ready application, as physiological functions are deeply integrated. In these organisms, the furthest extremities are physiologically linked to the central corpus of the individual. In this way, they are nearly opposite the concept of comprising semi-independent units. There is a critical disconnect, however, regarding this perception. Modules are traditionally thought of as “individuoids” or “complete physiological units,” compartments which package the full suite of physiological functions of an individual. But this does not need to be the case.

We can conceive of redundant organs in a subsystem, which might offer independence in function. So long as injury can be localised to a physiological system, one can envision a benefit conferred by redundancies. Unitisation can be within a single sub-system that ultimately affects the individual’s fitness. Albeit counterintuitive, there exists a practical translation of the conceptual model of modularity to highly physiologically integrated organisms (*i.e.*, traditional unitary organisms, such as higher vertebrates).

Injury to the physiology of integrated organisms is, almost by definition, of whole-organismal consequence. Importantly, integrated organisms typically relegate reproductive capacity to discrete sexual organs, and are thereby largely unamenable to regional reproductive isolation. This bauplan contrasts with that of plants, corals, and sponges, where both somatic and reproductive functions are carried out in unison. In those organisms, the death of a branch or sub-colony may not have important consequences. With a unitary animal, the loss of a subpart, such as an appendage or segment, may affect motility in a manner largely absent or irrelevant to plants and colonial invertebrates (Kozlowski 1973).

It is thus challenging to find a ready and competent gross analogue in most animal species. But as was recognised decades ago, and indicated above, the full or near-full range of individualistic properties exhibited by plant modules is not the qualification for modularity. The bauplan of the individual may not be the appropriate level of assessment (Tuomi and Mutikainen 1999; Harper 1980). Indeed, the axioms of modularity are multicellularity and multiple *redundant* units (Tuomi and Vuorisalo 1989). Thus, modularity is not necessarily incompatible with high bodily integration.

Archived information on the number of replicated organs across the tree of animals is surprisingly lacking. Tissues are often similar in organs across different species of animals and play a common functional role. We believe this to be a logical starting point for a systematic inquiry into the potential role of modularity in animal senescence, as posed by Finch (1990). Are any organs divided into major functional compartments that have functionally similar –if not equivalent– roles? Are they functionally related such that the impairment of one would not be indispensible to the individual’s survival?

We carried out an exhaustive survey classifying redundant components in 151 animal species. For the two systems – lymphatic and renal– where there appear to be conditions that lend themselves to functional redundancy, we investigate the types and scale of repeated elements in the system and how those are distributed among major groups of organisms. Table S3 of the online supplemental material contains a summary review of other organ systems. To test the effect of organ multiplicity on senescence, we used methods similar to those employed in plants (above). This involves running a series of phylogenetic generalised least square models to quantify the relationship between indices of organ multiples and mean life expectancy (*η*_e_; Table 1), maximum life expectancy (*η*_max_), and Keyfitz’ entropy (*H*).

#### 2.2.1 Immune system hierarchies

The lymphatic system is a strong candidate for a modular structure that might influence senescence partly because of the importance of immune integrity to survivorship (Nussey et al. 2013). More specifically, in certain species, the lymphatic system comprises numerous independent sub-units that are spatially discrete. Within species that have lymph nodes, we speculate that modularity corresponds with the number of lymph nodes in the body. Thus, species with a greater lymphatic modularity should have lower rates of senescence, as an extension of Finch’s thesis (1990).

Lymphatic systems of mammals span some twenty-fold variation in lymph node number, from 22 in mice to 450 in humans (Haley 2016) and several thousands in large ungulates (Nikles and Heath 1992). However, despite such a vast range of interspecific variation, reliable estimates of lymph node numbers across a large range of animal species do not exist. Indeed, for species lacking lymph nodes, other proxies for the degree of modularity require more finessing. Organs known as lymph hearts promote the movement of lymph through the lymph vascular network of certain species. They are found *in lieu* of lymph nodes in reptiles (Hedrick et al. 2013), amphibians (Hedrick et al. 2013), and some bird species (Hedrick et al. 2013; Budras, Hullinger, and Rautenfeld 1987). These structures can range from several to >200 in an individual (e.g. Caecilians, *Caecilia*; Kampmeier 1969).

Lymph heart function is not equivalent to that of lymph nodes. These organs do, however, share qualities in that they both influence immune system integrity and comprise discrete, highly numerated structures that mediate lymph. Independent of the above structures, temporary or reactive lymphatic structures develop in certain other species. These are variously termed aggregates, nodules, or patches, and participate in immune function. As such, they play a role in preventing tissue damage, disease, or dysfunction, and their continued functionality thereby reduces mortality. One of the complications with the lymphatic system in regard to isolating injury and compartmentalising damage is the functional integration of lymph nodes. Depending on the configuration, topology, and dynamics of lymph flow, regionalised isolation of injury may be infeasible.

#### 2.2.2. Renal hierarchies

The kidney presents a good example of a non-conventional morphological unit that would meet the discrete, functional unit criteria of modularity. Kidneys are physically separate and individually dispensible. As reflected in humans, loss of a kidney does not attend serious acute or long-term declines in the integrity of the renal system (Foley and Ibraim 2010; Wee, Tegzes, and Donker 1990 and Kasiske et al. 2013).

Most vertebrates possess two bilaterally positioned kidneys (Yokota et al. 2008). Among species, there is wide variation in both function and morphology of kidneys (Worthy and Williams 2002; Ortiz 2001; Beuchat 1996; Bentley 1971). In bears and cetaceans, for example, the kidney contains numerous lobes (*reniculation*) which exercise partial independence from one another (Maluf and Gassman 1998). The lobes comprise a physically integrated organ, but sub-regionalisation within this organ is high. This represents the highest degree of subdivision within our grading system because the kidney structure can be functionally divided into more than one-hundred compartments in some species (Ortiz 2001). Reniculated kidneys with high structural and functional division may offer more potential for localised damage and tissue death without functional injury to the renal system.

Kidney pathologies are diverse, but mostly internally mediated (Basile, Anderson, and Sutton 2012). Common drivers of kidney stress, injuiry, and death are high amino acid or nitrogenous waste loads, hypertension, and ischemia/hypoxia (Palm and Nordquist 2011). External factors such as blunt force trauma or kidney rupture more tightly akin to air embolism from the standpoint of acute, anatomically focused injury, are less common. It is unclear how subdivision affects the resiliency of renal physiology in response to different types of injury and stress.

While other species lack multi-compartmental renal organs, many species have multiple organs that participate in the management of wastes and ion balance. In marine fishes (Evans 2008, 2010), sharks (Shadwick, Farrell, and Brauner 2016), and crabs (Towle and Weihrauch 2001; Weihrauch and O’Donnel 2015), gills play an important role as an exchange site for ions between the organisms and their environments. In fact, these can be critical for nitrogenous waste excretion (see Wilkie 2002 and Wright and Wood 2012 for reviews). Many species of bony fishes are long lived, including dogfishes, paddlefishes, sturgeon, rock fishes, and eels (Patnaik, Mahapatro, Jena 1994). Similarly, a number of shark species are atypically long-lived (Hamady et al. 2014) and crustaceans, though particularly lacking in demographic studies, are popularly cited for their long lifespan (Vogt 2012). Despite this, there is little clade-level synthesis of senescence or negligible senescence for these groups (Patnaik, Mahapatro, Jena 1994; Vogt 2012). Attending the lack of research on senescence, links between specific anatomic similarities and longevity have not been directly explored, let alone interrogated in regard to renal anatomy.

Depressed osmoregulatory capacity resulting from injury to either the kidneys or the gills may be partially compensated for by having multiple osmoregulatory organs. Secondary osmoregulatory organs might also enable a higher osmoregulatory capacity and thereby tolerance for renal stress (Kültz 2015), which could both prevent injury and facilitate recovery (Reimschuessel 2001) from sub-lethal renal injury. In marine birds (Schmidt-Nielsen 1960) and some reptiles (Schmidt-Nielsen and Fange 1958), salt glands provide a mechanism to shed high salt concentrations. These glands could similarly enhance redundancy of function (Gutiérrez 2014; but see Babonis, Miller, and Evans 2011) without nitrogenous excretion. It is broadly recognised that birds are unusually long-lived (Hickey et al. 2012; Péron et al. 2010; Nussey et al. 2008; Monaghan et al. 2008; Ricklefs 2008), but research has conspicuously overlooked the role of anatomy in their longevity, and instead focused on cellular (Monaghan et al. 2008) and ecological explanations (Ricklefs 2008). Reptiles include a number of the longest-lived species, such as tuatara which can live some 90 years (Magalhaes and Costa 2009). Similar to birds, however, we see an inclination toward metabolic and cell-based explanations and an absence of attention given to organismal physiology and anatomy.

Along with gills, salt glands in accessory to kidneys constitute the second grade of centralisation in in renal function in animals (Fig 3c & d). They are ranked below reniculated kidneys because lobes of reniculation have the full suite of functional tools for renal control, whereas gills and salt glands are more limited. Gills in certain species of elasmobranchs (e.g. Wood, Pärt, and Wright 1995, Smith 1929) and salt-glands carry functional weight only as it relates to ionic homeostasis, not nitrogenous wastes control. Recorded variation tracks with environmental stresses, such as seawater ingestion (Worthy and Williams 2002) and hibernation (Ortiz 2001), and corresponds in part with acute and chronic kidney injury susceptibility (Stenvinkel et al. 2018).

We note that two-component systems, such as bilateral kidneys, raise special questions under the framework we describe here. These cases elevate questions about alternative explanations for the second organ that may not reflect an adaptation for functional redundancy. There are two particularly important scenarios: alternative selective pressures and non-adaptive tendencies.

As highlighted by Ewers and colleagues (2007), the function of two eyes is an adept example of how non-redundant traits can drive redundant features. As such, functional benefits unrelated to redundancy can confound the evolution of multiple organs and their potential role in modularity as it relates to senescence. Separate ears present another example of a highly specialised alternative benefit. More generally, symmetrical bauplans may favor bilateral, paired structures. This may be particularly likely where symmetrical division of gross anatomy shapes a developmental or functional pattern in the organism. A candidate for this might be the bilaterial lungs found in many terrestrial vertebrates. This issue has not been the subject of focused research.

Evolutionarily, the configuration of developmental control genes may influence the development of accessory organs (Oakley 2007). There is currently a dearth of research on the relative tendency for singular versus multiple organ duplication resulting from genetic mutations. Regardless, the cost of having multiple organs is likely to factor into whether organs are likely to fix into a population by genetic drift. If we assume that costs are commensurate with organ number, lower numbers of organs would have a higher likelihood of persisting in a population before being removed. Therefore, one should attribute functional purpose to singe-duplicate organs with a degree of skepticism.

## 3 ANALYSIS OF MODULARITY IN PLANTS AND ANIMALS

Our analysis supports Finch’s hypothesis that modularity influences senescence in plants, but support appears mixed across animals (Fig 3). We found seven relationships between modular traits and measures of senescence. The strongest relationships were found between between cumulative vessel redundancy and the pace of senescence (e.g., cumulative vessel redundancy × reproductive window (F=3.9; R^2^=0.41; Fig 3b) and between vessel density and the shape of senescence (F=4.2; R^2^=0.08; Fig 3a). However, most of the relationships we tested within plants were not statistically significant (Table S4).

In animals, there is evidence both supporting and refuting Finch’s hypothesis. Our tests support the hypothesis that modularity lengthens the pace of senescence in animals (Fig 3; Table S5). Species with lymphatic organs with greater subdivision into repeat components have significantly higher maximum lifespans (95% C.I. [36.4, 73.4 years]) than those with more centralised lymphatic anatomies (95% C.I. [19.0, 59.2 years]). On the other hand, evidence refuting Finch’s hypothesis was somewhat stronger. Lymphatic anatomies with greater subdivision into repeat components presented significantly lower mean life expectancies than those with lower subdivision (95% C.I. [4.1, 12.1 years] *vs.* 95% C.I. [7.9, 15.7 years]). When analysing renal anatomies, higher modular indices correlated with lower maximum lifespans (95% C.I. [0.3, 13.6 years]) and we found no correlation between modularity index and mean life expectancy. The renal modularity index was significantly negatively correlated with the shape of senescence (95% C.I [0.008, 0.084]). Other relationships were insignificant (Table S5).

## 4 DISCUSSION AND FUTURE DIRECTIONS

In this study, we tested Finch’s hypothesis (1990) that more modular organisms should be more likely to live longer because of a decoupling of local injury from individual mortality. We situated Finch’s hypothesis in a broad, comparative framework to look at a complete spectrum of degrees of physiological integration within the context of the most recent research on senescence (Vaupel et al. 2004; Jones et al. 2014; Cohen 2018). To test our hypotheses in a fully comparative framework, we used functional traits to explore plant senescence and propose measures of modularity in unitary animals, a group of organisms thus far neglected in the comparative study of modularity. We found support for Finch’s hypothesis in plants, but mixed support in animals.

Our results suggest that, in plants, modularity may in fact prevent or moderate the potential for localised injuries to depress demographic rates (*i.e.,* reproduction and survival). Modularity may decouple localised injuries from individual fitness. Our work offers a general test of this hypothesis using functional traits and demographic data, and highlights the need for further work to uncover underlying processes.

We tap a similar angle for two animal traits for which our results are less conclusive. In the two physiological systems we studied in animals, the effect of modularised systems on the pace and shape of senescence may be limited because of (i) the types of mortality factors they interact with (Williams et al. 2006; Caswell 2007; Baudisch and Vaupel 2010); (ii) the potentially confounding influence of other unitary physiological systems to which they are linked (physiological causality, and thus relationships, are not well understood; Nussey et al. 2013 and Rickefs 2008); or (iii) because their preliminary classification of subdivision does not best represent functional divisions (trait-based approaches based on organ morphology are few).

### Plants

In plants, the examination of this empirical support suggests that modularity can affect senescence in a wide variety of ways. In particular, modularity could affect senescence by changing average mortality throughout the lifecycle of organisms, by decreasing age-specific mortality rates. Modularity could also affect senescence by increasing the likelihood of reaching maturity and the reproductive window with age (Caswell 2007).

We found a positive correlation between modularity in plants and the pace and shape of senescence. This suggests that the degree of modularity may affect the life cycle by interacting with age-independent and age-specific mortality rates. A consistent pattern in the slowed pace of senescence between modularity and clonality supports the view that there is similarity in demographic process. Extended lifespans that we report for modular plants adds to, and generalises from, a tradition of studies that relate clonal forms to longevity (Orive 1995; de Witte and Stöcklin 2010; Sköld and Obst 2011). Modularity can be thought of, in limited part, as a downscaled, localized, and weaker form of semi-clonality (White 1979). This trait enables a larger, composite organisms to access a new ecological and evolutionary space that is inaccessible to less unitary plants (Mackie 1986). Today, the spectrum of modularity adds a complex layer to the picture of demography, by grading a key distinction regarding senescence between clonal and non-clonal species. Unlike clonality, which lies at the high end of the physiological independence spectrum, many other modular species tend to have an intermediate degree of physiological integration. Modular species therefore present a valuable opportunity to explore senescence and the mechanism which relieve it.

Moreover, the measures of modularity that correlate with measures of senescence further support the adaptive value of hydraulic integration as suggested by Schenk and colleagues (2008). Though promising, we cannot, with the data available at present, attribute the general relationships in our results to the causality of air embolism and related injury. Moreover, the potential adaptive aspects of modularity may not speak to the origins of its evolution, or if it is incidental to another adaptive feature in plants.

The correlations between stem density/cumulative vessel redundancy and mean life expectancy (a measure of the pace of senescence; Fig. 2) suggest that modularity decreases the average mortality rate. However, the correlation between the same measures of modularity and a less strong shape of senescence (Keyfitz’ entropy; Keyfitz 1977) could reflect an alleviation of age-specific risk factors. Our initial expectation was that modularity would not influence the shape of senescence, but for a specific caveat related to mutation accumulation theory (*sensu* Medawar 1952).

Our expectation implied that the effect of modularity would not change based on the age of an organism. A correlation between modularity and the shape of senescence suggests that modularity can affect age-specific mortality factors. Such decrease in age-specific mortality rates could be a direct or indirect effect of modularity. Our measures of modularity –including stem density and cumulative vessel redundancy– might plausibly decrease age-specific mortality directly. This direct decrease could occur because the risk of stem cavitation should increase with the height, and therefore age, of individual plants (Koch et al. 2004). However, other modularity traits could decrease age-specific mortality rates indirectly. For example, epicormic branching should increase the capacity for long-term growth (Ishii et al. 2001) and therefore fertility (Ishii et al. 2007) of trees. In this case, modularity could decrease age-specific mortality rates indirectly, by increasing size.

### Animals

In animals, our analyses partly support the hypothesis that modularity should decrease the pace of senescence in animals. In particular, we found that immune anatomies with greater organ system subdivision have significantly higher mean lifespans than those with higher centralisation. However, we found an opposite (though weaker) trend when looking at mean lifespan.

The nexus of immune anatomy and immune function is an area that would benefit from further study, particularly because of the role of localization and metastatic processes. Moreover, there is a dearth of studies on the functional property differences between lymph nodes and lymph hearts (Hendrick et al. 2013). In particular, there is little available information on interspecific lymph node variation. But despite having effectively no information on lymph node variation, the category has a narrow distribution of rates of senescence (*H)* and mean life expectancies (*η_e_*). *H-values* reflect that not only most organisms exhibited similarly lengthed lifespans, but that mortality patterns were similar among species. Lymph nodes are almost exclusive to class Mammalia, for which the overall number of samples in the data base is not high – and thus this may be partly an artifact of sample size. This topic remains of high interest because of growing interest in immunosenescence and how lymphatics intermediate immune integraity comparative physiology of lymph (Aw, Silva, and Palmer 2007).

In animals, the rate of senescence was not consistently predicted by our index of modularity. One possible explanation for these results may reside in the way we classified basal vertebrate and invertebrate systems as “centralised” or non-modular. There is also latitude to interpret which sets and subsets of organs are appropriate for comparative analysis, which could prove material to the results. Given the extreme range of anatomies and distributed nature of available data on anatomical biometrics, there is ample room, apt timing, and increasing value (Griffith et al. 2016; Weiss and Ray 2019) in a more refined approach to defining functional and anatomical trait groups for comparison.

An opposite trend between modularity and longevity in renal anatomy is notable. In addition to caveats noted at the outset of this section, this result may be explained partly by variation in the renal stresses experienced by the organisms evaluated. For example, renal stress is greater for species which ingest high amounts of salt and for those that exchange ions with hypertonic environments (Hildebrandt 2001). Similar influences may be responsible for the unexpected pattern we identified for age-specific mortality differences in renal anatomy.

In both plants and animals, there was a lack of many significant relationships between measures of modularity and measures of senescence. The significant relationships might reflect spurious correlations or true, reproducible patterns. Prior knowledge suggests that our significant correlations likely reflect reproducible patterns, at least in the case of plants. We expect these patterns to be reproducible because our significant correlations include measures of modularity that are considered adaptive (Schenk et al. 2008). Future studies should clarify the strength of the evidence here presented, especially because the sample sizes of freely available demographic and functional trait data are increasing exponentially with time (Enquist et al. 2009, Kattge et al. 2011, Salguero-Gómez et al. 2015, 2016, Iversen et al. 2017).

In species where reproduction is limited by actuarial senescence, we would expect modularity to change the shape of reproductive senescence (Ricklefs 2008; Lemaître and Gaillard 2017). Specifically, modularity that promotes, directly or indirectly, the attainment of large organismal size should increase age-class dependent resproduction (Stearns 1992). A relative increase in an organism’s maximum organismal size at first reproduction should do the same (Lacey 1986). We do not analytically test this in our review due to data constraints. We note, however, that contrary to our *a priori* expectation, the literature provides evidence of reproductive senescence in clonal species (e.g. Ally et al. 2010) and actuarial senescence in asexual species (Martinez and Levinton 1992).

We do not test reproductive senescence in this monograph because there are more involved procedures required to ascertain changes in reproductive patterns. These are not supported by the current data available, but may be in the near future. Nevertheless, we predict that modularity could affect reproductive senescence if it facilitates the attainment of large sizes, to which fertility is often proportional (Weiner et al. 2009). In addition, at large sizes, the patterns of selection for fertility and mortality can become decoupled from age (Caswell and Salguero-Gómez 2013; Roper, Capdevila, and Salguero-Gómez 2019). Forthcoming work lays a conceptual foundation for statistically decomposing fertility patterns similarly to how we approach actuarial patterns here (Baudisch and Stott 2019). This approach uses pace and shape metrics in a slightly different application, offering a promising next step to this subject.

Our analyses of plant and animal data shows that greater degrees of modularity correlate with the pace and shape of senescence, at least in plants. Recent developments in research on hydraulic integration provide strong inference for why we would expect to see the patterns we report. Despite this, one cannot eliminate the possibility that correlated, yet to be identified benefits are responsible for driving this pattern. Moreover, current mechanisms do not necessarily give insight to the evolutionary origins of the trait, which will require addition study. These results offer a foundation from which to work and enhance the urgency for this line of research.

## Future directions

Our findings and review point to three avenues of potentially fruitful future research:

### Age-specific mortality factors influenced by modularity

We have long understood that modularity, at least in the form of clones, could relieve mortality forces for the greater individual (Williams 1957; Cook 1979; Salguero-Gómez et al. 2013). That less-physiologically independent modules could achieve a similar outcome and play a role in determining the shape of senescence, is a more recent area of thought (Vaupel et al. 2004). While we are left with more questions than answers in our results, our finding that modularity may act on age-specific mortality rates suggests value in a research agenda focusing on age-specific mortality factors as a fitness benefit of modularity.

### Role of dysregulation and physiological modules in the endocrine system

The field would benefit from additional investigation into the role of dysregulation and physiological modules in the endocrine system. A growing body of research describes a close connection between endocrine dysregulation and ageing in humans (Li et al. 2015; Cohen et al. 2013; López-Otín et al. 2013). This work provides an important and potentially widely-applicable mechanism for ageing, explaining patterns of physiological senescence, based on the analysis of biomarkers. Dysregulation research advances are at the forefront of understanding intermediating factors in physiological declines. Theory in this area suggests that dysregulation does not necessarily rely upon the three prevailing theories for the evolution of senescence (Cohen et al. 2016). Rather, this phenomenon is likely linked to emergent properties of complex systems. Understanding the role of this mechanism, its origination, and its interactions with related theory –including the modularity inferface– will be a major step in senescence research.

### Specific limitation on the fitness advantage of modularity

Ageing biologists would benefit from a research programme that probes the specific limitations on the fitness advantage conferred by modularity. Welch and Waxman (2003) offer a strong example of the type of work to be probed at this interface from an evolutionary angle. We have found reduced mortality rates associated with highly modularised architectures in plants. A now relevant question is whether there are costs and specific factors which curtail such a benefit to plants and animals. Some candidates for inquiry include environmental stresses that could be exacerbated by branched morphologies (in climes where snow, wind, cold temperatures, or other physical stresses exert routine injury), energetic costs of higher surface area (bark-to-vascular tissue), vulnerability to disease, and light competition factors.

For decades, senescence was believed to be a universal phenomenon. Evidence now exists that the ageing process is not inevitable in all species. Mechanisms remain elusive, but this paper takes a modest step forward in testing Finch’s prediction of one mechanism based on the modular architecture of plants and animals. Future research is needed to more deeply explore relevant modular traits and their influence on specific mortality factors in plant and animal species.

## Supporting information

Online Supplement. Species Tables.

## ACKNOWLEDGEMENTS

This research was supported by an Australian Research Council grant (DE140100505) and a NERC IRF (R/142195-11-1) to RSG. The COMPADRE & COMADRE databases have been supported by the Max Planck Institute for Demographic Research. Some trait data were obtained from the TRY database, which is curated at the Max Planck Institute for Biogeochemistry with support from DIVERSITAS, IGBP, the Global Land Project, NERC QUEST, the Royal Botanic Gardens Kew SID, FRB, and GIS “Climat, Environnement et Société” France. We also thank the FRED and BIEN initiatives for their full open-access support, and B. Maitner and B. Enquist for help during data processing on the latter. The authors declare no conflict of interest.

## AUTHORS’ CONTRIBUTIONS

RSG conceived the ideas and designed methodology with input from CB and AC. RSG quantified life history traits. AC collated information on plant functional traits. CB obtained indicators of modularity in animals. RSG and CB ran the phylogenetic analyses. CB led the writing of the manuscript with input from AC and RSG.

## DATA ACCESSIBILITY

Data used in this manuscript will be deposit open access in DRYAD when the work is accepted. The demographic information is permanently, open-access archived in www.compadre-db.org.

