## Supplementary material for "Testing Finch’s hypothesis: the role of organismal modularity on the escape from actuarial senescence": Online Supplement. Species Tables.

**Connor Bernard, Aldo Compagnoni & Roberto Salguero-Gómez**

**Table S1. List of species in this review**

**Table S2. List of traits linked to modularity in plants**

**Table S3.** List of traits linked to modularity in animals

**Table S1.** List of species (138 plants and 151 animal species) used in the review for which functional traits and demographic information were obtained from various resources to explore the morphological correlates of senescence. The bibliographic references correspond to sources of demographic information from COMPADRE & COMADRE.

Plants:

| **Species** | **Family** | **Authors** | **Journal** | **DOI.ISBN** | **Year**  **Published** |
| --- | --- | --- | --- | --- | --- |
| *Arenaria serpyllifolia* | Caryophyllaceae | Dostál | J Veg Sci | 10.1658/1100-9233(2007)18[91:PDOAIP]2.0.CO;2 | 2007 |
| *Cerastium fontanum* | Caryophyllaceae | Burns; Pardini; Schutzenhofer; Chung; Seidler; Knight | Ecology | 10.1890/12-1310.1 | 2013 |
| *Chenopodium album* | Amaranthaceae | Davis | Weed Sci | 10.1890/1051-0761(2006)016[2399:DMISOB]2.0.CO;2 | 2006 |
| *Erigeron canadensis* | Compositae | Bullock; White; Prudhomme; Tansey; Perea; Hooftman | J Ecol | 10.1111/j.1365-2745.2011.01910.x | 2011 |
| *Oryza sativa* | Poaceae | Vidotto; Ferrero; Ducco | Weed Res | 10.1046/j.1365-3180.2001.00244.x | 2001 |
| *Achillea millefolium* | Compositae | Fréville; Silvertown | Plant Ecol | 10.1007/s11258-004-0017-1 | 2005 |
| *Agropyron cristatum* | Poaceae | Hansen; Wilson | J Appl Ecol | 10.1111/j.1365-2664.2006.01145.x | 2006 |
| *Elymus repens* | Poaceae | Mortimer | Book | NA | 1983 |
| *Alliaria petiolata* | Brassicaceae | Meekins; McCarthy | Am Midl Nat | 10.1674/0003-0031(2002)147[0256:EOPDOT]2.0.CO;2 | 2002 |
| *Alliaria petiolata* | Brassicaceae | Drayton; Primack | Biol Inv | NA | 1999 |
| *Alliaria petiolata* | Brassicaceae | Burns; Pardini; Schutzenhofer; Chung; Seidler; Knight | Ecology | 10.1890/12-1310.1 | 2013 |
| *Alliaria petiolata* | Brassicaceae | Evans; Davis; Raghu; Ragavendran; Landis; Schemske | Ecol Appl | 10.1890/11-1291.1 | 2012 |
| *Anthoxanthum odoratum* | Poaceae | Fréville; Silvertown | Plant Ecol | 10.1007/s11258-004-0017-1 | 2005 |
| *Rytidosperma caespitosum* | Poaceae | Williams; Wills; Janes; Vander Schoor; Newton; Hovenden | New Phytol | 10.1111/j.1469-8137.2007.02170.x | 2007 |
| *Carex bigelowii* | Cyperaceae | Carlsson; Callaghan | Oikos | 10.2307/3544870 | 1991 |
| *Chamaecrista fasciculata* | Leguminosae | Stephens; Tye; Quintana-Ascencio | Popul Ecol | 10.1007/s10144-014-0438-1 | 2014 |
| *Cleome droserifolia* | Cleomaceae | Hegazy | J Arid Env | NA | 1990 |
| *Daucus carota* | Apiaceae | Verkaar; Schenkeveld | New Phyto | 10.1111/j.1469-8137.1984.tb04155.x | 1984 |
| *Geranium sylvaticum* | Geraniaceae | Ramula; Toivonen; Mutikainen | Int J Plant Sci | 10.1086/512040 | 2007 |
| *Hypochaeris radicata* | Compositae | Jongejans; de Kroon | J Ecol | 10.1111/j.1365-2745.2005.01003.x | 2005 |
| *Lotus corniculatus* | Leguminosae | Emery; Beuselinck; English | New Phyto | 10.1046/j.1469-8137.1999.00540.x | 1999 |
| *Molinia caerulea* | Poaceae | Jacquemyn; Brys; Neubert | Ecol Appl | 10.1890/04-1762 | 2005 |
| *Picris hieracioides* | Compositae | Klemow; Raynal | J Ecol | 10.2307/2259775 | 1985 |
| *Podophyllum peltatum* | Berberidaceae | Sohn; Policansky | Ecology | 10.2307/1935088 | 1977 |
| *Ranunculus acris* | Ranunculaceae | Sarukhan; Harper | J Ecol | 10.2307/2258643 | 1973 |
| *Rubus saxatilis* | Rosaceae | Eriksson | Ecol Research | 10.1007/BF02348412 | 1994 |
| *Rubus ursinus* | Rosaceae | Lambrecht-McDowell; Radosevich | Biol Inv | 10.1007/s10530-004-0870-9 | 2005 |
| *Sporobolus heterolepis* | Poaceae | Dalgleish; Kula; Hartnett; Sandercook | Am J Bot | 10.3732/ajb.2007277 | 2008 |
| *Taraxacum campylodes* | Compositae | Vavrek; McGraw; Yang | J Ecol | 10.2307/2960501 | 1997 |
| *Taraxacum campylodes* | Compositae | Burns; Pardini; Schutzenhofer; Chung; Seidler; Knight | Ecology | 10.1890/12-1310.1 | 2013 |
| *Themeda triandra* | Poaceae | O'Connor; Pickett | J Appl Ecol | 10.2307/2404276 | 1992 |
| *Themeda triandra* | Poaceae | Williams; Wills; Janes; Vander Schoor; Newton; Hovenden | New Phytol | 10.1111/j.1469-8137.2007.02170.x | 2007 |
| *Trifolium pratense* | Leguminosae | Fréville; Silvertown | Plant Ecol | 10.1007/s11258-004-0017-1 | 2005 |
| *Trollius europaeus* | Ranunculaceae | Lemke; Salguero-Gomez | Popul Ecol | 10.1007/s10144-015-0519-9 | 2015 |
| *Euterpe precatoria* | Arecaceae | Zuidema | Book | NA | 2000 |
| *Euterpe precatoria* | Arecaceae | Otárola; Avalos | Am J Bot | 10.3732/ajb.1400089 | 2014 |
| *Iriartea deltoidea* | Arecaceae | Pinard | Biotrop | 10.2307/2388974 | 1993 |
| *Rhopalostylis sapida* | Arecaceae | Enright; Watson | New Zealand J Bot | 10.1080/0028825X.1992.10412883 | 1992 |
| *Acacia aneura* | Leguminosae | Rosenberg; Boland; Tiver; Watson | Internet | NA | 2005 |
| *Acacia victoriae* | Leguminosae | Grice; Westoby; Torpy | Austral Ecol | 10.1111/j.1442-9993.1994.tb01537.x | 1994 |
| *Betula nana* | Betulaceae | Ebert; Ebert | Vegetatio | 10.1007/BF00042253 | 1989 |
| *Calluna vulgaris* | Ericaceae | Scandrett; Gimmingham | Vegetatio | 10.1007/BF00036515 | 1989 |
| *Cornus florida* | Cornaceae | Vejdani | PhD thesis | NA | 2006 |
| *Hydrangea paniculata* | Hydrangeaceae | Hara; Kanno; Hirabuki; Takehara | J Veg Sci | 10.1111/j.1654-1103.2004.tb02286.x | 2004 |
| *Lantana camara* | Verbenaceae | Osunkoya; Perrett; Fernando; Clark; Raghu | Popul Ecol | 10.1007/s10144-013-0364-7 | 2013 |
| *Lindera benzoin* | Lauraceae | Cipollini; Wallace-Senft; Whigham | J Ecol | 10.2307/2261269 | 1994 |
| *Lonicera maackii* | Caprifoliaceae | Burns; Pardini; Schutzenhofer; Chung; Seidler; Knight | Ecology | 10.1890/12-1310.1 | 2013 |
| *Miconia prasina* | Melastomataceae | Pascarella; Alde; Zimmerman | Biotrop | 10.1111/j.1744-7429.2006.00220.x | 2007 |
| *Rhododendron maximum* | Ericaceae | McGraw | Am J Bot | 10.2307/2444780 | 1989 |
| *Rosa canina* | Rosaceae | Burns; Pardini; Schutzenhofer; Chung; Seidler; Knight | Ecology | 10.1890/12-1310.1 | 2013 |
| *Spartium junceum* | Fabaceae | Stevens; Latimer | Glob Change Biol | 10.1111/gcb.12824 | 2014 |
| *Viburnum furcatum* | Adoxaceae | Hara; Kanno; Hirabuki; Takehara | J Veg Sci | 10.1111/j.1654-1103.2004.tb02286.x | 2004 |
| *Abies concolor* | Pinaceae | van Mantgem; Stephenson | J Ecol | 0.1111/j.1365-2745.2005.01007.x | 2005 |
| *Abies magnifica* | Pinaceae | van Mantgem; Stephenson | J Ecol | 0.1111/j.1365-2745.2005.01007.x | 2005 |
| *Abies sachalinensis* | Pinaceae | Kubota | Eco Research | 10.1007/BF02523604 | 1997 |
| *Acer palmatum* | Sapindaceae | Tanaka; Shibata; Masaki; Iida; Niiyama; Abe; Kominomi; Nokashizuka | J Veg Sci | 10.3170/2007-8-18342 | 2008 |
| *Acer pictum* | Sapindaceae | Tanaka; Shibata; Masaki; Iida; Niiyama; Abe; Kominomi; Nokashizuka | J Veg Sci | 10.3170/2007-8-18342 | 2008 |
| *Acer rufinerve* | Sapindaceae | Tanaka; Shibata; Masaki; Iida; Niiyama; Abe; Kominami; Nokashizuka | J Veg Sci | 10.3170/2007-8-18342 | 2008 |
| *Acer saccharum* | Sapindaceae | Lin; Augspurger | Forest Ecol Manag | 10.1016/j.foreco.2008.02.040 | 2008 |
| *Adansonia digitata* | Malvaceae | Venter; Witkowski | Forest Ecol Manag | 10.1016/j.foreco.2013.04.013 | 2013 |
| *Aesculus turbinata* | Sapindaceae | Kaneko; Takada; Kawano | Plant Spp Biol | 10.1046/j.1442-1984.1999.00007.x | 1999 |
| *Ailanthus altissima* | Simaroubaceae | Burns; Pardini; Schutzenhofer; Chung; Seidler; Knight | Ecology | 10.1890/12-1310.1 | 2013 |
| *Anisoptera laevis* | Dipterocarpaceae | Yamada; Yamada; Okuda; Fletcher | Oecologia | 10.1007/s00442-012-2529-z | 2013 |
| *Aquilaria malaccensis* | Thymelaeaceae | Soehartono; Newton | Biol Cons | 10.1016/S0006-3207(00)00089-6 | 2001 |
| *Araucaria araucana* | Araucariaceae | Bekessy; Newton; Fox; Lara et al. | Book | NA | 2004 |
| *Araucaria cunninghamii* | Araucariaceae | Enright; Ogden | Aust J Ecol | 10.1111/j.1442-9993.1979.tb01195.x | 1979 |
| *Araucaria hunsteinii* | Araucariaceae | Enright | Aust J Ecol | 10.1111/j.1442-9993.1982.tb01304.x | 1982 |
| *Avicennia germinans* | Acanthaceae | López-Hoffman; Ackerly; Anten; Denoyer;Ramos | J Ecol | 10.1111/j.1365-2745.2007.01298.x | 2007 |
| *Avicennia marina* | Acanthaceae | Burns; Ogden | Aust J Ecol | 10.1111/j.1442-9993.1985.tb00874.x | 1985 |
| *Bertholletia excelsa* | Lecythidaceae | Zuidema; Boot | J Trop Ecol | 10.1017/S0266467402002018 | 2002 |
| *Brosimum alicastrum* | Moraceae | Peters | PhD thesis | NA | 1989 |
| *Callitris columellaris* | Cupressaceae | Price; Bowman | J Biogeog | 10.2307/2846032 | 1994 |
| *Calocedrus decurrens* | Cupressaceae | van Mantgem; Stephenson | J Ecol | 10.1111/j.1365-2745.2005.01007.x | 2005 |
| *Calocedrus macrolepis* | Cupressaceae | Chien; Zuidema; Nghia | Popul Ecol | 10.1007/s10144-008-0079-3 | 2008 |
| *Camellia japonica* | Theaceae | Shimatani; Kubota; Araki; Aikawa; Manobe | Plant Spp Biol | 10.1111/j.1442-1984.2007.00190.x | 2007 |
| *Carapa guianensis* | Meliaceae | Klimas; Cropper Jr.; Kainer; Wadt | Ecol Model | 10.1016/j.ecolmodel.2012.07.022 | 2012 |
| *Castanea dentata* | Fagaceae | Davelos; Jarosz | J Ecol | 10.1111/j.0022-0477.2004.00907.x | 2004 |
| *Cecropia obtusifolia* | Urticaceae | Alvarez-Buylla | Am Nat | 10.1086/285599 | 1994 |
| *Cedrela odorata* | Meliaceae | Zuidema; Brienen; During; Guneralp | Am Nat | 10.1086/605981 | 2009 |
| *Chlorocardium rodiei* | Lauraceae | ter Steege; Boot; Brouwer; Hammond; Vanderhout; Jetten; Khan; Polak; Raaimakers; Zagt | Ecol Appl | 10.2307/2269341 | 1995 |
| *Choerospondias axillaris* | Anacardiaceae | Brodie; Helmy; Brockelman; Maron | Ecol Appl / Ecol | 10.1890/08-0111.1 | 2009 |
| *Dacrydium elatum* | Podocarpaceae | Chien; Zuidema; Nghia | Popul Ecol | 10.1007/s10144-008-0079-3 | 2008 |
| *Dicorynia guianensis* | Leguminosae | Picard; Mortier; Chagneau | Ecol Model | 10.1016/j.ecolmodel.2010.06.010 | 2010 |
| *Dicymbe altsonii* | Leguminosae | Zagt; Boot | PhD thesis | NA | 1997 |
| *Entandrophragma cylindricum* | Meliaceae | Picard; Yalibanda; Namkosserena; Baya | Forest Ecol Manag | 10.1016/j.foreco.2008.02.041 | 2008 |
| *Eperua falcata* | Leguminosae | Chagneau; Mortier; Picard | J R Stat Soc C | 10.1111/j.1467-9876.2008.00657.x | 2009 |
| *Euptelea pleiosperma* | Eupteleaceae | He; Wang; Franklin; Jiang | Biol Cons | 10.1016/j.biocon.2013.03.011 | 2013 |
| *Fagus crenata* | Fagaceae | Nakashizuka | J Veg Sci | NA | 1991 |
| *Fagus grandifolia* | Fagaceae | Batista; Platt; Macchiavelli | Ecology | 10.2307/176863 | 1998 |
| *Fagus grandifolia* | Fagaceae | Batista; Platt; Macchiavelli | Ecology | 10.2307/176863 | 1998 |
| *Fagus sylvatica* | Fagaceae | López; Ortuño; Martin; Fullano | Ann For Sci | 10.1051/forest:2007037 | 2007 |
| *Grias peruviana* | Lecythidaceae | Peters | Book | NA | 1991 |
| *Guaiacum sanctum* | Zygophyllaceae | CITES | Plants Committee | NA | 2008 |
| *Khaya senegalensis* | Meliaceae | Gaoue; Ticktin | Cons Biol | 10.1111/j.1523-1739.2009.01345.x | 2010 |
| *Khaya senegalensis* | Meliaceae | Gaoue; Ticktin | Cons Biol | 10.1111/j.1523-1739.2009.01345.x | 2010 |
| *Khaya senegalensis* | Meliaceae | Gaoue;  Horvitz; Ticktin; Steiner; Tuljapurkar | J Ecol | 10.1111/1365-2745.12140 | 2013 |
| *Melaleuca viridiflora* | Myrtaceae | Crowley; Garnett; Shephard | Aust Ecol | 10.1111/j.1442-9993.2008.01921.x | 2009 |
| *Microberlinia bisulcata* | Leguminosae | Norghauer; Newbery | Ecol Monog | 10.1890/10-2268.1 | 2011 |
| *Nothofagus fusca* | Nothofagaceae | Enright; Ogden | Aust J Ecol | 10.1111/j.1442-9993.1979.tb01195.x | 1979 |
| *Ocotea usambarensis* | Lauraceae | Stas; Langbroek; Bitariho; Sheil; Zuidema | Afr J Ecol | NA | 2016 |
| *Oxandra asbeckii* | Annonaceae | Chagneau; Mortier; Picard | J R Stat Soc C | 10.1111/j.1467-9876.2008.00657.x | 2009 |
| *Parkinsonia aculeata* | Leguminosae | Raghu; Wilson; Dhileepan | Aust J Ent | 10.1111/j.1440-6055.2006.00556.x | 2006 |
| *Pentaclethra macroloba* | Leguminosae | Hartshorn | PhD thesis | NA | 1972 |
| *Phyllanthus emblica* | Phyllanthaceae | Sinha; Brault | Biodivers Conserv | 10.1007/s10531-004-0827-4 | 2005 |
| *Phyllanthus emblica* | Phyllanthaceae | Ticktin; Ganesan; Paramesha; Setty | J Appl Ecol | 10.1111/j.1365-2664.2012.02156.x | 2012 |
| *Phyllanthus emblica* | Phyllanthaceae | Ellis; Williams; Lesica; Bell; Bierzychudek; Bowles; Crone; Doak; Ehrlén; Ellis-Adam; McEachern; Ganesan; Latham; Luijten; Kaye; Knight; Menges; Morris; Den Nijs; Oostermeijer; Quintana-Ascencio; Shelly; Stanley; Thorpe; Ticktin; Valverde; Weekley | Ecology | 10.1890/11-1052.1 | 2012 |
| *Picea jezoensis* | Pinaceae | Kubota | Eco Research | 10.1007/BF02523604 | 1997 |
| *Pinus jeffreyi* | Pinaceae | van Mantgem; Stephenson | J Ecol | 10.1111/j.1365-2745.2005.01007.x | 2005 |
| *Pinus fenzeliana* | Pinaceae | Chien; Zuidema; Nghia | Popul Ecol | 10.1007/s10144-008-0079-3 | 2008 |
| *Pinus lambertiana* | Pinaceae | van Mantgem; Stephenson | J Ecol | 10.1111/j.1365-2745.2005.01007.x | 2005 |
| *Pinus lambertiana* | Pinaceae | Maloney; Vogler; Eckert; Jensen; Neale | Forest Ecol Manag | 10.1016/j.foreco.2011.05.011 | 2011 |
| *Manilkara zapota* | Sapotaceae | Cruz-Rodriguez; López-Villavicencio; Valverde | J Trop Ecol | 10.1017/S0266467408005713 | 2009 |
| *Pinus nigra* | Pinaceae | Buckley; Brockerhoff; Langer; Ledgard; North; Rees | J Appl Ecol | 10.1111/j.1365-2664.2005.01100.x | 2005 |
| *Pinus palustris* | Pinaceae | Platt; Evans; Rathbun | Am Nat | 10.1086/284803 | 1988 |
| *Pinus ponderosa* | Pinaceae | van Mantgem; Stephenson | J Ecol | 10.1111/j.1365-2664.2005.01100.x | 2005 |
| *Pinus strobus* | Pinaceae | Münzbergová; Hadincová; Wild; Kindlmannová | PLoS ONE | 10.1371/journal.pone.0056953 | 2013 |
| *Pinus radiata* | Pinaceae | Reynolds | PhD thesis | NA | 2015 |
| *Pinus sylvestris* | Pinaceae | Usher | Biom | 10.2307/2401258 | 1966 |
| *Prioria copaifera* | Leguminosae | Condit | Forest Ecol Manag | 10.1016/0378-1127(93)90045-O | 1993 |
| *Prosopis flexuosa* | Fabaceae | Aschero; Morris; Vázquez; Alvarez; Villagra | For Ecol Manag | 10.1016/j.foreco.2016.03.028 | 2016 |
| *Prosopis glandulosa* | Leguminosae | Golubov; Mandujano; Franco; Montaña; Eguiarte; Lopez-Portillo | J Ecol | 10.1046/j.1365-2745.1999.00420.x | 1999 |
| *Prunus africana* | Rosaceae | Stewart | PhD thesis | NA | 2001 |
| *Prunus serotina* | Rosaceae | Sebert-Cuvillier; Paccaut; Chabrerie; Endels; Goubet; Decoq | Ecol Model | 10.1016/j.ecolmodel.2006.09.005 | 2007 |
| *Psidium guajava* | Myrtaceae | Somarriba | Agrofor Syst | 10.1007/BF02344742 | 1988 |
| *Pterocarpus angolensis* | Leguminosae | Desmet; Shackleton; Ronbinson | S African J Bot | NA | 1996 |
| *Rhizophora mangle* | Rhizophoraceae | López-Hoffman; Ackerly; Anten; DeNoyer;Ramos | J Ecol | 10.1111/j.1365-2745.2007.01298.x | 2007 |
| *Rhododendron ponticum* | Ericaceae | Salguero-Gómez | MSc thesis | NA | 2004 |
| *Rhododendron ponticum* | Ericaceae | Travis; Harris; Park; Bullock | MEE | 10.1111/j.2041-210X.2011.00104.x | 2011 |
| *Roupala montana* | Proteaceae | Hoffmann | Ecology | 10.2307/177080 | 1999 |
| *Sapium sebiferum* | Euphorbiaceae | Renne | PhD thesis | NA | 2001 |
| *Scaphium macropodum* | Malvaceae | Yamada; Zuidema; Itoh; et al. | J Ecol | 10.1111/j.1365-2745.2006.01209.x | 2007 |
| *Scaphium macropodum* | Malvaceae | Le | NA | 978-90-393-5778-1 | 2012 |
| *Sclerocarya birrea* | Anacardiaceae | Emanuel | Forest Ecol Manag | 10.1016/j.foreco.2005.03.066 | 2005 |
| *Sequoia sempervirens* | Cupressaceae | Namkoong; Roberds | Am Nat | 10.1086/282913 | 1974 |
| *Sequoia sempervirens* | Cupressaceae | Bosch | Science | 10.1126/science.172.3981.345 | 1971 |
| *Shorea acuminata* | Dipterocarpaceae | Yamada; Yamada; Okuda; Fletcher | Oecologia | 10.1007/s00442-012-2529-z | 2013 |
| *Shorea bracteolata* | Dipterocarpaceae | Yamada; Yamada; Okuda; Fletcher | Oecologia | 10.1007/s00442-012-2529-z | 2013 |
| *Shorea leprosula* | Dipterocarpaceae | Visser; Jongejans; van Breugel; Zuidema; Chen; Kassim; de Kroon | J Ecol | 10.1111/j.1365-2745.2011.01825.x | 2011 |
| *Shorea maxwelliana* | Dipterocarpaceae | Yamada; Yamada; Okuda; Fletcher | Oecologia | 10.1007/s00442-012-2529-z | 2013 |
| *Shorea ovalis* | Dipterocarpaceae | Yamada; Yamada; Okuda; Fletcher | Oecologia | 10.1007/s00442-012-2529-z | 2013 |
| *Stryphnodendron microstachyum* | Leguminosae | Hartshorn | PhD thesis | NA | 1972 |
| *Styrax obassis* | Styracaceae | Abe; Nokashizuka; Tanoka | J Veg Sci | 10.2307/3237044 | 1998 |
| *Swietenia macrophylla* | Meliaceae | Verwer; Peña-Claros; van der Staak; Ohlson-Kiehn; Sterck | J Appl Ecol | 10.1111/j.1365-2664.2008.01564.x | 2008 |
| *Syzygium jambos* | Myrtaceae | Brown; Spector; Wu | J Appl Ecol | 10.1111/j.1365-2664.2008.01550.x | 2008 |
| *Taxus brevifolia* | Taxaceae | Busing; Spies | USDAResearchNate | NA | 1995 |
| *Tetraberlinia bifoliolata* | Leguminosae | Norghauer; Newbery | Ecol Monog | 10.1890/10-2268.1 | 2011 |
| *Tsuga canadensis* | Pinaceae | Lamar; McGraw | Forest Ecol Manag | 10.1016/j.foreco.2005.02.056 | 2005 |
| *Vatica mangachapoi* | Dipterocarpaceae | Hu; Wang | Acta Ecol Sinica | NA | 1988 |
| *Vochysia ferruginea* | Vochysiaceae | Boucher; Mallono | Forest Ecol Manag | 10.1016/S0378-1127(96)03890-X | 1997 |
| *Vouacapoua americana* | Leguminosae | Chagneau; Mortier; Picard | J R Stat Soc C | 10.1111/j.1467-9876.2008.00657.x | 2009 |
| *Ziziphus jujuba* | Rhamnaceae | Zull; Laws; Cacho | Environ Modell Softw | 10.1016/j.envsoft.2015.10.026 | 2015 |

Animals

| **Species** | **Family** | **Authors** | **Journal** | **DOI.ISBN** | **Year**  **Publication** |
| --- | --- | --- | --- | --- | --- |
| *Abax parallelepipedus* | Carabidae | Pichancourt; Burel; Auger | C R Biol | 10.1016/j.crvi.2005.09.009 | 2006 |
| *Accipiter cooperii* | Accipitridae | Schumaker; Ernst; White; Baker; Haggerty | Ecol Appl | 10.1890/02-5010 | 2004 |
| *Accipiter gentilis* | Accipitridae | Schumaker; Ernst; White; Baker; Haggerty | Ecol Appl | 10.1890/02-5010 | 2004 |
| *Acinonyx jubatus* | Felidae | Crooks; Sanjayan; Doak | Biol Cons | 10.1046/j.1523-1739.1998.97054.x | 1998 |
| *Acipenser fulvescens* | Acipenseridae | Velez-Espino; Koops | N Am J Fish Manag | 10.1577/M08-034.1 | 2009 |
| *Acipenser fulvescens* | Acipenseridae | Velez-Espino; Koops | N Am J Fish Manag | 10.1577/M08-034.1 | 2009 |
| *Acipenser transmontanus* | Acipenseridae | Velez-Espino; Koops | Ecol Model | 10.1016/j.ecolmodel.2012.09.022 | 2012 |
| *Acropora cervicornis* | Acroporidae | Mercado-Molina; Ruiz-Diaz; Pérez; Rodríguez-Barreras; Sabat | Coral Reef | 10.1007/s00338-015-1341-8) | 2015 |
| *Acropora downingi* | Acroporidae | Riegl; Purkis | Glob Change Biol | 10.1111/gcb.13014 | 2015 |
| *Acropora hyacinthus* | Acroporidae | Tanner | J Exp Mar Biol & Ecol | 10.1016/S0022-0981(97)00024-5 | 1997 |
| *Acyrthosiphon pisum* | Aphididae | Hamda; Jevtic; Laskowski | Ecotox | 10.1007/s10646-012-0904-5 | 2012 |
| *Adamussium colbecki* | Pectinidae | Ripley; Caswell | Popul Ecol | 10.1007/s10144-008-0075-7 | 2008 |
| *Aedes albopictus* | Culicidae | Hanly; Haase | J Med Entomol | 10.1093/jme/tjw021 | 2016 |
| *Aepyceros melampus* | Bovidae | Spinage | Ecology | 10.2307/1934778 | 1972 |
| *Agaricia agaricites* | Agariciidae | Hughes; Tanner | Ecology | 10.1890/0012-9658(2000)081[2250:RFLHAL]2.0.CO;2 | 2000 |
| *Agelaius phoeniceus* | Icteridae | Blackwell; Huszar; Linz; Dolbeer | J Wildlife Manag | 10.2307/3802689 | 2003 |
| *Ailuropoda melanoleuca* | Ursidae | Carter; Ackleh; Leonard; Wang | Ecol Model | 10.1016/S0304-3800(99)00145-3 | 1999 |
| *Alcedo atthis* | Alcedinidae | Gleason; Nacci | Hum & Ecol Risk Assess | 10.1080/20018091094835 | 2001 |
| *Alces alces* | Cervidae | Ballard; Whitman; Reed | Wildlife Monogr | 10.2307/3830713 | 1991 |
| *Alcyonium sp.* | Alcyoniidae | McFadden | Ecology | 10.2307/1940983 | 1991 |
| *Alligator mississippiensis* | Alligatoridae | Tucker | Book | 978-0-949324-89-4 | 2001 |
| *Amazona vittata* | Psittacidae | Beissinger; Wunderle; Meyers; Saether; Engen | Ecol Monogr | 10.1890/07-0018.1 | 2008 |
| *Ambloplites rupestris* | Centrarchidae | Peoples | Master Thesis | NA | 2010 |
| *Ambystoma mexicanum* | Ambystomatidae | Zambrano; Vega; Herrera; Prado; Reynoso | Anim Conserv | 10.1111/j.1469-1795.2007.00105.x | 2007 |
| *Ammocrypta pellucida* | Percidae | Velez-Espino; Koops | Ecol Model | 10.1016/j.ecolmodel.2012.09.022 | 2012 |
| *Ammodramus savannarum* | Emberizidae | Hovick; Miller | Land Ecol | 10.1007/s10980-013-9896-7 | 2013 |
| *Ampelisca abdita* | Ampeliscidae | Kuhn; Munns; Serbst; Edwards; Cantwell; Gleason; Pelletier; Berry | Env Tox & Chem | 10.1002/etc.5620210425 | 2002 |
| *Amphiascus tenuiremis* | Diosaccidae | Chandler; Cary; Bejarano; Pender; Ferry | Environ Sci Technol | 10.1021/es049654o | 2004 |
| *Amphimedon compressa* | Niphatidae | Mercado-Molina; Sabat; Yoshioka | J Exp Mar Biol & Ecol | 10.1016/j.jembe.2011.07.018 | 2011 |
| *Amphiprion percula* | Pomacentridae | Buston; Garcia | J Fish Biol | 10.1111/j.1095-8649.2007.01445.x | 2007 |
| *Amphispiza belli subsp. belli* | Emberizidae | Naujokaitis-Lewis; Curtis; Arcese; Rosenfeld | Conserv Biol | 10.1111/j.1523-1739.2008.01066.x | 2009 |
| *Anarhynchus frontalis* | Charadriidae | Cruz; Pech; Seddon; Cleland; Nelson; Sanders; Maloney | Biol Cons | 10.1016/j.biocon.2013.09.006 | 2013 |
| *Anas fulvigula* | Anatidae | Rigby; Haukos | Southeast Nat | 10.1656/058.013.s505 | 2014 |
| *Anas laysanensis* | Anatidae | Reynolds; Weiser; Jamieson; Hatfield | J Wild Manag | 10.1002/jwmg.582 | 2013 |
| *Anas platyrhynchos* | Anatidae | Hoekman; Mills; Howerter; Devries; Ball | J Wild Manag | 10.2307/3803153 | 2002 |
| *Anaxyrus boreas* | Bufonidae | Biek; Funk; Maxell; Mills | Conserv Biol | 10.1046/j.1523-1739.2002.00433.x | 2002 |
| *Anser anser* | Anatidae | Klok; van Turnhout; Willems; Voslamber; Ebbinge; Schekkerman | Anim Biol | 10.1163/157075610X523260 | 2010 |
| *Antechinus agilis* | Dasyuridae | Lindenmayer; Lacy | Biol Cons | 10.1016/S0006-3207(01)00134-3 | 2002 |
| *Anthropoides paradiseus* | Gruidae | Altwegg; Anderson | Funct Ecol | 10.1111/j.1365-2435.2009.01563.x | 2009 |
| *Apalone mutica* | Trionychidae | Zimmer-Shaffer; Briggler; Millspaugh | Chel Cons Biol | 10.2744/CCB-1109.1 | 2014 |
| *Apalone spinifera* | Trionychidae | Zimmer-Shaffer; Briggler; Millspaugh | Chel Cons Biol | 10.2744/CCB-1109.1 | 2014 |
| *Aquila fasciata* | Accipitridae | Chevallier; Hernandez-Matias; Real; Vincent-Martin; Ravayrol; Basnard | J Appl Ecol | 10.1111/1365-2664.12476 | 2015 |
| *Arctica islandica* | Arcticidae | Ripley; Caswell | Popul Ecol | 10.1007/s10144-008-0075-7 | 2008 |
| *Arctodiaptomus salinus* | Diaptomidae | Jiménez-Melero; Gilbert; Guerrero | Fresh Biol | 10.3354/meps10377 | 2013 |
| *Assa darlingtoni* | Myobatrachidae | Keith; Mahony; Hines; Elith; Regan; Baumgartner; Hunter; Heard; Mitchell; Parris; Penman; Scheele; Simpson; Tingley; Tracy; West; Akcakaya | Conserv Biol | 10.1111/cobi.12234 | 2014 |
| *Astroblepus ubidiai* | Astroblepidae | Velez-Espino | Ecol Freshwater Fish | 10.1111/j.1600-0633.2005.00084.x | 2005 |
| *Aythya affinis* | Anatidae | Koons | Avian Cons and Ecol | NA | 2006 |
| *Bonasa umbellus* | Phasianidae | Blomberg; Tefft; Reed; McWilliams | J Wildlife Manage | 10.1002/jwmg.278 | 2011 |
| *Bos frontalis* | Bovidae | Ahrestani; Iyer; Heitkönig; Prins | Mammal Rev | 10.1111/j.1365-2907.2010.00166.x | 2011 |
| *Bos primigenius* | Bovidae | Lesnoff; Corniaux; Hiernaux | Ecol Model | 10.1016/j.ecolmodel.2012.02.018 | 2012 |
| *Bostrychia hagedash* | Threskiornithidae | Duckworth; Altwegg; Harebottle | J Orni | 10.1007/s10336-011-0758-2 | 2011 |
| *Botrylloides violaceus* | Styelidae | Grey | Oecologia | 10.1007/s00442-011-1931-2 | 2011 |
| *Botryllus schlosseri* | Styelidae | Cockrell; Sorte | J Exp Mar Bio & Ecol | 10.1016/j.jembe.2012.11.010 | 2013 |
| *Brachyrhaphis rhabdophora* | Poeciliidae | Johnson; Zuniga-Vega | Ecology | 10.1890/07-1672.1 | 2009 |
| *Brachyteles hypoxanthus* | Atelidae | Morris; Altmann; Brockman; Cords; Fedigan; Pusey; Stoinski; Bronikowski; Alberts; Strier | Am Nat | 10.1086/657443 | 2011 |
| *Brevicoryne brassicae* | Aphididae | Ciblis-Stewart; Sandercock; McCornack | Entomol Exp Appl | 10.1111/eea.12325 | 2015 |
| *Bubo virginianus* | Strigidae | Schumaker; Ernst; White; Baker; Haggerty | Ecol Appl | 10.1890/02-5010 | 2004 |
| *Buteo buteo* | Accipitridae | Meyer; Meyer; Francisco; Holder; Verdonck | PLOS One | 10.1371/journal.pone.0147189 | 2016 |
| *Buteo jamaicensis* | Accipitridae | Schumaker; Ernst; White; Baker; Haggerty | Ecol Appl | 10.1890/02-5010 | 2004 |
| *Buteo lineatus* | Accipitridae | Lawler; Schumaker | Environ Monit Assess | 10.1023/B:EMAS.0000016881.12925.b1 | 2004 |
| *Buteo solitarius* | Accipitridae | Klavitter; Marzluff; Vekasy | J Wild Manage | 10.2307/3803072 | 2003 |
| *Caenorhabditis elegans* | Rhabditidae | Li; Ju; Liao; Liao | Ecotoxicology | 10.1007/s10646-014-1267-x) | 2014 |
| *Caiman crocodilus* | Alligatoridae | Tucker | Book | 978-0-949324-89-4 | 2001 |
| *Calidris temminckii* | Scolopacidae | Koivula; Pakanen; Rönkä; Belda | Avian Biol | 10.1111/j.0908-8857.2008.04189.x | 2008 |
| *Callinectes sapidus* | Portunidae | Miller | Estuaries | 10.2307/1353238 | 2001 |
| *Callorhinus ursinus* | Otariidae | Barlow; Boneng | Mar Mammal Sci | 10.1111/j.1748-7692.1991.tb00550.x | 1991 |
| *Callospermophilus lateralis* | Sciuridae | Hostetler; Kneip; Van Vuren; Oli | PLoS1 | 10.1371/jourNAl.pone.0034379 | 2012 |
| *Calyptorhynchus lathami* | Psittacidae | Harris; Fordham; Mooney; Pedler; Araújo; Paton; Stead; Watts; Akçakaya; Brook | J Appl Ecol | 10.1111/j.1365-2664.2012.02163.x | 2012 |
| *Camelus dromedarius* | Camelidae | Pflaumer | JSM | NA | 2013 |
| *Campylorhynchus brunneicapillus subsp. sandiegensis* | Troglodytidae | Conlisk; Motheral; Chung; Wisinski; Endress | Biol Cons | 10.1016/j.biocon.2014.04.010 | 2014 |
| *Canis latrans* | Canidae | Schumaker; Ernst; White; Baker; Haggerty | Ecol Appl | 10.1890/02-5010 | 2004 |
| *Canis lupus* | Canidae | Chapron; Legendre; Ferrière; Clobert; Haight | C R Biol | 10.1016/S1631-0691(03)00148-3 | 2003 |
| *Canis lupus familiaris* | Canidae | Makenov; Bekova | Urban Ecosyst | 10.1007/s11252-016-0566-9 | 2016 |
| *Capitella sp.* | Capitellidae | Hansen; Forbes; Forbes | Funct Ecol | 10.1046/j.1365-2435.1999.00299.x | 1999 |
| *Capra ibex* | Bovidae | Gaillard; Yoccoz | Ecology | 10.1890/02-0409 | 2003 |
| *Cardisoma guanhumi* | Gecarcinidae | Rodriguez-Fourquet | PhD thesis | NA | 2004 |
| *Caretta caretta* | Cheloniidae | Crouse; Crowder; Caswell | Ecology | 10.2307/1939225 | 1987 |
| *Castor canadensis* | Castoridae | Payne | J Wildlife Manage | 10.2307/3808459 | 1984 |
| *Catostomus catostomus* | Catostomidae | Velez-Espino; Koops | Ecol Model | 10.1016/j.ecolmodel.2012.09.022 | 2012 |
| *Catostomus commersoni* | Catostomidae | Miller; Tietge; McMaster; Munkittrick; Xia; Griesmer; Ankley | Env Tox | 10.1002/etc.2972 | 2015 |
| *Catostomus platyrhynchus* | Catostomidae | Young; Koops | Fish & Oceans Can | NA | 2013a |
| *Cebus capucinus* | Cebidae | Morris; Altmann; Brockman; Cords; Fedigan; Pusey; Stoinski; Bronikowski; Alberts; Strier | Am Nat | 10.1086/657443 | 2011 |
| *Centrocercus minimus* | Stercorariidae | Davis; Hooten; Phillips; Doherty | Ecology and Evolution | 10.1002/ece3.1290 | 2014 |
| *Centrocercus urophasianus* | Phasianidae | Johnson | Cons Biol | 10.1046/j.1523-1739.1999.97284.x | 1999 |
| *Cephaloleia fenestrata* | Chrysomelidae | Johnson; Horvitz | Ecology | 10.1890/04-0974 | 2005 |
| *Cercopithecus mitis* | Cercopithecidae | Morris; Altmann; Brockman; Cords; Fedigan; Pusey; Stoinski; Bronikowski; Alberts; Strier | Am Nat | 10.1086/657443 | 2011 |
| *Certhia americana* | Certhiidae | Wintle; Bekessy; Venier; Pearce; Chisholm | Cons Biol | 10.1111/j.1523-1739.2005.00276.x | 2005 |
| *Cervus canadensis subsp. nelsoni* | Cervidae | Clark | PhD Thesis | NA | 2014 |
| *Cervus elaphus* | Cervidae | Benton; Grant; Clutton-Brock | Evol Ecol | 10.1007/BF01237655 | 1995 |
| *Chelodina expansa* | Chelidae | Spencer; Thomson | Cons Biol | 10.1111/j.1523-1739.2005.00487.x | 2005 |
| *Chelonia mydas* | Cheloniidae | Chaloupka | Ecol Model | 10.1016/S0304-3800(01)00433-1 | 2002 |
| *Chelydra serpentina* | Chelydridae | Congdon; Dunham; van Loben Sels | Am Zool | 10.1093/icb/34.3.397 | 1994 |
| *Chen caerulescens* | Anatidae | Cooch; Rockwell; Brault | Ecol Monog | 10.1890/0012-9615(2001)071[0377:RAODRT]2.0.CO;2 | 2001 |
| *Chrosomus oreas* | Cyprinidae | Peoples | Master Thesis | NA | 2010 |
| *Chrysemys picta* | Emydidae | Wilbur | Ecology | 10.2307/1935300 | 1975 |
| *Cicindela ohlone* | Carabidae | Cornelisse; Bennett; Letourneau | PLOS One | 10.1371/jourNAl.pone.0071005 | 2013 |
| *Ciconia ciconia* | Ciconiidae | Schaub; Pradel; Lebreton | Biol Cons | 10.1016/j.biocon.2003.11.002 | 2004 |
| *Cistothorus palustris* | Troglodytidae | Schumaker; Ernst; White; Baker; Haggerty | Ecol Appl | 10.1890/02-5010 | 2004 |
| *Clemmys guttata* | Emydidae | Enneson; Litzgus | Biol Cons | 10.1016/j.biocon.2008.04.001 | 2008 |
| *Clethrionomys rufocanus* | Muridae | Yoccoz; Nakata; Stenseth; Saitoh | Res Popul Ecol | 10.1007/BF02765226 | 1998 |
| *Clethrionomys sp.* | Muridae | Row; Wilson; Murray | J Anim Ecol | 10.1111/1365-2656.12179 | 2014 |
| *Clinocottus analis* | Cottidae | Davis; Levin | Mar Ecol Prog Series | 10.3354/meps234229 | 2002 |
| *Clinocottus globiceps* | Cottidae | Pfister | Ecology | 10.1007/bf02027951 | 1996 |
| *Clinostomus elongatus* | Cyprinidae | Velez-Espino; Koops | Ecol Model | 10.1016/j.ecolmodel.2012.09.022 | 2012 |
| *Clinostomus funduloides* | Cyprinidae | Peoples | Master Thesis | NA | 2010 |
| *Colias alexandra* | Pieridae | Hayes | Oecologia | 10.1007/BF00349187 | 1981 |
| *Coragyps atratus* | Cathartidae | Blackwell; Avery; Watts; Lowney | J Wild Manage | 10.2193/2006-146 | 2007 |
| *Coregonus huntsmani* | Salmonidae | Velez-Espino; Koops | Ecol Model | 10.1016/j.ecolmodel.2012.09.022 | 2012 |
| *Coregonus reighardi* | Salmonidae | Velez-Espino; Koops | Ecol Model | 10.1016/j.ecolmodel.2012.09.022 | 2012 |
| *Coregonus zenithicus* | Salmonidae | Velez-Espino; Koops | Ecol Model | 10.1016/j.ecolmodel.2012.09.022 | 2012 |
| *Cornu aspersa* | Helicidae | Laskowski; Hopkin | Ecotox Environ Safe | 10.1006/eesa.1996.0045 | 1996 |
| *Cottus bairdi* | Cottidae | Peoples | Master Thesis | NA | 2010 |
| *Cottus confusus* | Cottidae | Velez-Espino; Koops | Ecol Model | 10.1016/j.ecolmodel.2012.09.022 | 2012 |
| *Cottus sp.* | Cottidae | Velez-Espino; Koops | Ecol Model | 10.1016/j.ecolmodel.2012.09.022 | 2012 |
| *Crassostrea virginica* | Ostreidae | Puckett | PhD Thesis | NA | 2013 |
| *Crocodylus acutus* | Crocodylidae | Tucker | Book | NA | 2001 |
| *Crocodylus johnsoni* | Crocodylidae | Tucker | Book | 978-0-949324-89-4 | 2001 |
| *Crocodylus niloticus* | Crocodylidae | Hutton | PhD | NA | 1984 |
| *Cryptobranchus alleganiensis subsp. alleganiensis* | Cryptobranchidae | Unger; Sutton; Williams | J Nat Cons | 10.1016/j.jnc.2013.06.002 | 2013 |
| *Cryptophis nigrescens* | Elapidae | Webb; Brook; Shine | Ecol Res | 10.1046/j.1440-1703.2002.00463.x | 2002 |
| *Cyphastrea microphthalma* | Faviidae | Riegl; Purkis | Glob Change Biol | 10.1111/gcb.13014 | 2015 |
| *Cyprinodon diabolis* | Cyprinodontidae | Beissinger | PeerJ | 10.7717/peerj.549 | 2014 |
| *Cyprinus carpio* | Cyprinidae | Stratford; Pollino; Brown | Environ Modell Softw | 10.1016/j.envsoft.2016.02.009 | 2016 |
| *Daphnia magna* | Daphniidae | Duchet; Coutellec; Franquet; Lagneau; Lagadic | Ecotoxic | 10.1007/s10646-010-0507-y | 2011 |
| *Daphnia pulex* | Daphniidae | Duchet; Coutellec; Franquet; Lagneau; Lagadic | Ecotoxic | 10.1007/s10646-010-0507-y | 2010 |
| *Dasyatis violacea* | Dasyatidae | Mollet; Cailliet | Mar Freshwater Res | 10.1071/MF01083 | 2002 |
| *Dendragapus obscurus* | Phasianidae | Schumaker; Ernst; White; Baker; Haggerty | Ecol Appl | 10.1890/02-5010 | 2004 |
| *Diadema antillarum* | Diadematidae | Rodríguez-Barreras; Pérez; Mercado-Molina; Sabat | Estuar Marin Shelf S | 10.1016/j.ecss.2015.06.021 | 2015 |
| *Diceros bicornis* | Rhinocerotidae | Brodie; Muntifering; Hearn; Loutit; Loutit; Brell; Uri-Khob; Leader-Williams; Preez | Anim Conserv | 10.1111/j.1469-1795.2010.00434.x | 2011 |
| *Didelphis aurita* | Didelphidae | Ferreira; Kajin; Vieira; Zangrandi; Cerqueira; Gentile | Mammal Biol | 10.1016/j.mambio.2013.03.002 | 2013 |
| *Diomedea exulans* | Diomedeidae | Koons | PhD Thesis | NA | 2005 |
| *Diploria strigosa* | Faviidae | Edmunds | Mar Ecol Prog Ser | 10.3354/meps08595 | 2010 |
| *Dipsastrea pallida* | Merulinidae | Riegl; Purkis | Glob Change Biol | 10.1111/gcb.13014 | 2015 |
| *Drymarchon couperi* | Colubridae | Hyslop; Stevenson; Macey; Carlile; Jenkins; Hostetler; Oli | Popul Ecol | 10.1007/s10144-011-0292-3 | 2011 |
| *Dryocopus pileatus* | Picidae | Schumaker; Ernst; White; Baker; Haggerty | Ecol Appl | 10.1890/02-5010 | 2004 |
| *Trichechus manatus latirostris* | Trichechidae | Heinsohn; Lacy; Lindenmayer; Marsh; Kwan; Lawler | Anim Conserv | 10.1017/S1367943004001593 | 2004 |
| *Eidolon helvum* | Pteropodidae | Hayman; McCrea; Restif; Suu-Ire; Fooks; Wood; Cunningham; Rowcliffe | J Mamm | 10.1017/S0950268812000167 | 2012 |
| *Eisenia fetida* | Lumbricidae | Santadino; Coviella; Momo | Water Air Soil Pollut | 10.1007/s11270-014-2207-3 | 2014 |
| *Elephas maximus* | Elephantidae | Chelliah; Bukka; Sukumar | Biol Cons | 10.1016/j.biocon.2013.05.008 | 2013 |
| *Emydura macquarii* | Chelidae | Spencer; Thomson | Cons Biol | 10.1111/j.1523-1739.2005.00487.x | 2005 |
| *Enhydra lutris* | Mustelidae | Gerber; Tinker; Doak; Estes; Jessup | Ecol Appl | 10.1890/03-5006 | 2004 |
| *Entosphenus macrostomus* | Petromyzontidae | Velez-Espino; Koops | Ecol Model | 10.1016/j.ecolmodel.2012.09.022 | 2012 |
| *Epidalea calamita* | Bufonidae | Di Minin; Griffiths | Ecography | 10.1111/j.1600-0587.2010.06263.x | 2011 |
| *Epinephelus morio* | Serranidae | Fujiwara; Zhou | Can J Fish Aq Sci | 10.1139/cjfas-2012-0520 | 2013 |
| *Erimyzon sucetta* | Catostomidae | Velez-Espino; Koops | Ecol Model | 10.1016/j.ecolmodel.2012.09.022 | 2012 |
| *Esox lucius* | Esocidae | Edeline; Haugen; Weltzien; Claessen; Winfield; Stenseth; Vollestad | Proc R Soc B | 10.1098/rspb.2009.1724 | 2010 |
| *Etheostoma flabellare* | Percidae | Peoples | Master Thesis | NA | 2010 |
| *Eubalaena glacialis* | Balaenidae | Fujiwara; Caswell | Ecology | 10.2307/3072076 | 2002 |
| *Eulamprus tympanum* | Scincidae | Blomberg; Shine | Aust Ecol | 10.1046/j.1442-9993.2001.01120.x | 2001 |
| *Eumetopias jubatus* | Otariidae | Holmes; York | Cons Biol | 10.1111/j.1523-1739.2003.00191.x | 2003 |
| *Falco naumanni* | Falconidae | Hiraldo; Negro; Donazar; Gaona | J Appl Ecol | 10.2307/2404688 | 1996 |
| *Falco peregrinus* | Falconidae | Altwegg; Jenkins; Abadi | Ibis | 10.1111/ibi.12125 | 2013 |
| *Falco peregrinus subsp. anatum* | Falconidae | Deines; Peterson; Boeckner; Boyle; Keighley; Kogut; Lubben; Rebarber; Ryan; Tenhumberg; Townley; Tyre | Ecol Appl | 10.1890/06-1090.1 | 2007 |
| *Felis catus* | Felidae | Budke; Slater | J Appl Anim Welf Sci | 10.1080/10888700903163419 | 2009 |
| *Forpus passerinus* | Psittacidae | Sandercock; Beissinger | J Appl Stats | 10.1080/02664760120108818 | 2002 |
| *Fulmarus glacialis* | Procellariidae | Kerbiriou; Le Viol; Bonnet; Robert | Popul Ecol | 10.1007/s10144-012-0306-9 | 2012 |
| *Gasterosteus sp.* | Gasterosteidae | Velez-Espino; Koops | Ecol Model | 10.1016/j.ecolmodel.2012.09.022 | 2012 |
| *Gavia immer* | Gaviidae | Grear; Meyer; Cooley; Kuhn; Piper; Mitro; Vogel; Taylor; Kenow; Craig; Nacci | J Wild Manag | 10.2193/2008-093 | 2010 |
| *Gemma gemma* | Veneridae | Ripley; Caswell | Popul Ecol | 10.1007/s10144-008-0075-7 | 2008 |
| *Genypterus blacodes* | Ophidiidae | Gonzales-Olivares; Aranguiz-Acuna; Ramos-Jiliberto; Rojas-Palma | Fish Res | 10.1016/j.fishres.2008.11.006 | 2009 |
| *Geocrinia alba* | Myobatrachidae | Conroy; Brook | Pop Ecol | 10.1007/s10144-003-0145-9 | 2003 |
| *Geocrinia vitellina* | Myobatrachidae | Conroy; Brook | Pop Ecol | 10.1007/s10144-003-0145-9 | 2003 |
| *Geukensia demissa* | Mytilidae | Ripley; Caswell | Popul Ecol | 10.1007/s10144-008-0075-7 | 2008 |
| *Goniastrea aspera* | Faviidae | Babcock | Ecol Monog | 10.2307/2937107 | 1991 |
| *Goniastrea favulus* | Faviidae | Babcock | Ecol Monog | 10.2307/2937107 | 1991 |
| *Gopherus agassizii* | Testudinidae | Doak; Kareiva; Klepetka | Ecol Appl | 10.2307/1941949 | 1994 |
| *Gorgonia ventalina* | Gorgoniidae | Sabat; Toledo-Hernández | J Marin Biol | 10.1155/2015/987060 | 2015 |
| *Gorilla beringei* | Hominidae | Morris; Altmann; Brockman; Cords; Fedigan; Pusey; Stoinski; Bronikowski; Alberts; Strier | Am Nat | 10.1086/657443 | 2011 |
| *Gulo gulo* | Mustelidae | Carroll; Noss; Paquet; Schumaker | Ecol Appl | 10.1890/02-5195 | 2003 |
| *Gyps coprotheres* | Accipitridae | Monadjem; Wolter; Neser; Kane | Anim Conserv | 10.1111/acv.12054 | 2013 |
| *Haematopus ostralegus* | Haematopodidae | Klok; Roodbergen; Hemerik | Anim Biol | 10.1163/157075609X417143 | 2009 |
| *Haliaeetus albicilla* | Accipitridae | Krüger; Grünkorn; Struwe-Juhl | Biol Cons | 10.1016/j.biocon.2009.12.010 | 2010 |
| *Halichoerus grypus* | Phocidae | Harwood | J Appl Ecol | 10.2307/2402601 | 1978 |
| *Haliotis corrugata* | Haliotidae | Button; Rogers-Bennett | Mar Ecol Prog Ser | 10.3354/meps09094 | 2011 |
| *Haliotis laevigata* | Haliotidae | Fordham; Mellin; Russell; Akcakaya; Bradshaw; Aiello-Lammens; Caley; Connell; Mayfield; Shepherd; Brook | GCB | 10.1111/gcb.12289 | 2013 |
| *Haliotis rufescens* | Haliotidae | Rogers-Bennett; Leaf | Ecol Appl | 10.1890/04-1688 | 2006 |
| *Haliotis sorenseni* | Haliotidae | Rogers-Bennett; Leaf | Ecol Appl | 10.1890/04-1688 | 2006 |
| *Helioseris cucullata* | Agariciidae | Hughes; Tanner | Ecology | 10.1890/0012-9658(2000)081[2250:RFLHAL]2.0.CO;2 | 2000 |
| *Hemitragus jemlahicus* | Bovidae | Caughley | Ecology | 10.2307/1935638 | 1966 |
| *Himantopus novaezelandiae* | Recurvirostridae | Cruz; Pech; Seddon; Cleland; Nelson; Sanders; Maloney | Biol Cons | 10.1016/j.biocon.2013.09.005 | 2013 |
| *Hippocamelus bisulcus* | Cervidae | Corti; Wittmer; Festa-Bianchet | J Mamm | 10.1644/09-MAMM-A-047.1 | 2010 |
| *Hirundo rustica* | Hirundinidae | Grüebler; Korner-Nievergelt; Naef-Daenzer | Ecol & Evol | 10.1002/ece3.984 | 2014 |
| *Homo sapiens sapiens* | Hominidae | Keyfitz; Flieger | Book | 0-226-43237-8 | 1990 |
| *Hoplocephalus bungaroides* | Elapidae | Webb; Brook; Shine | Ecol Res | 10.1046/j.1440-1703.2002.00463.x | 2002 |
| *Huso huso* | Acipenseridae | Doukakis; Babcock; Pikitch; Sharov; Baimukhanov; Erbulekov; Bokova; Nimatov | Cons Biol | 10.1111/j.1523-1739.2010.01458.x | 2010 |
| *Hybognathus argyritis* | Cyprinidae | Velez-Espino; Koops | Ecol Model | 10.1016/j.ecolmodel.2012.09.022 | 2012 |
| *Hypseleotris klunzingeri* | Eleotridae | Yen; Bond; Shenton; Spring; Mac Nally | J Appl Ecol | 10.1111/1365-2664.12074 | 2013 |
| *Hystrix refossa* | Hystricidae | Monchot; Fernandez; Gaillard | J Arch Sci | 10.1016/j.jas.2012.04.037 | 2012 |
| *Isurus oxyrinchus* | Lamnidae | Tsai; Sun; Punt; Liu | ICES J Mar Sci | 10.1093/icesjms/fsu056 | 2014 |
| *Jynx torquilla* | Picidae | Schaub; Reichlin; Abadi; Kéry; Jenni; Arlettaz | Oecologia | 10.1007/s00442-011-2070-5 | 2012 |
| *Kinosternon flavescens* | Kinosternidae | Iverson | Herpetologica | 10.2307/1447430 | 1991 |
| *Kinosternon integrum* | Kinosternidae | Macip-Ríos; Brauer-Robleda; Zúñiga-Vega; Casas-Andreu | Herp J | NA | 2011 |
| *Kinosternon subrubrum* | Kinosternidae | Frazer; Gibbons; Greene | Ecology | 10.2307/1941572 | 1991 |
| *Kobus ellipsiprymnus* | Bovidae | Van Sickle; Atwell; Craig | J Wildlife Manage | 10.2307/3801764 | 1987 |
| *Lacerta agilis* | Lacertidae | Berglind | Ecol Bull | 10.2307/20113253 | 2000 |
| *Lagopus leucura* | Phasianidae | Wilson; Martin | BMC Ecol | 10.1186/1472-6785-12-9 | 2012 |
| *Lagopus muta* | Phasianidae | Wilson; Martin | BMC Ecol | 10.1186/1472-6785-12-9 | 2012 |
| *Lagopus muta subsp. japonica* | Phasianidae | Suzuki; Kobayashi; Nakamura; Takasu | Wildlife Biol | 10.2981/13-021 | 2013 |
| *Lagothrix lagotricha* | Atelidae | Defler | Book | 978-1-4939-0697-0 | 2014 |
| *Lampetra richardsoni* | Petromyzontidae | Velez-Espino; Koops | Ecol Model | 10.1016/j.ecolmodel.2012.09.022 | 2012 |
| *Lasaea rubra* | Lasaeidae | Ripley; Caswell | Popul Ecol | 10.1007/s10144-008-0075-7 | 2008 |
| *Lemmus lemmus* | Muridae | Row; Wilson; Murray | J Anim Ecol | 10.1111/1365-2656.12179 | 2014 |
| *Leopardus pardalis* | Felidae | Haines; Tewes; Laack; Grant; Young | Biol Cons | 10.1016/j.biocon.2005.06.032 | 2005 |
| *Lepeophtheirus salmonis* | Caligidae | Groner; Gettinby; Stormoen; Revie; Cox | PLOS One | 10.1371/jourNAl.pone.0088465 | 2014 |
| *Lepetodrilus fucensis* | Lepetodrilidae | Kelly; Metaxas | Mar Ecol Pro Ser | 10.3354/meps08442 | 2010 |
| *Lepisosteus oculatus* | Lepisosteidae | Velez-Espino; Koops | Ecol Model | 10.1016/j.ecolmodel.2012.09.022 | 2012 |
| *Leptogorgia virgulata* | Gorgoniidae | Gotelli | Ecology | 10.1046/j.1461-0248.2000.00138.x | 1991 |
| *Lepus americanus* | Leporidae | Meslow; Keith | J Wildlife Manage | 10.2307/3799557 | 1968 |
| *Lepus europaeus* | Leporidae | Marboutin; Peroux | J Appl Ecol | 10.2307/2404820 | 1995 |
| *Lichenostomus melanops subsp. cassidix* | Meliphagidae | Baxter; McCarthy; Possingham; Menkhorst; McLean | Cons Biol | 10.1111/j.1523-1739.2006.00378.x | 2006 |
| *Lissarca miliaris* | Philobryidae | Ripley; Caswell | Popul Ecol | 10.1007/s10144-008-0075-7 | 2008 |
| *Lissarca notorcadensis* | Philobryidae | Ripley; Caswell | Popul Ecol | 10.1007/s10144-008-0075-7 | 2008 |
| *Lontra canadensis* | Mustelidae | Gorman; McMillan; Erb; Deperno; Martin | Am Midl Nat | 10.1674/0003-0031(2008)159[98:SACMOA]2.0.CO;2 | 2008 |
| *Loxodonta africana* | Elephantidae | Chelliah; Bukka; Sukumar | Biol Cons | 10.1016/j.biocon.2013.05.008 | 2013 |
| *Lucanus miwai* | Lucanidae | Huang | J Asia-Pac Entomol | 10.1016/j.aspen.2014.03.009 | 2014 |
| *Lycalopex culpaeus* | Canidae | Novaro; Funes; Walker | J Appl Ecol | 10.1111/j.1365-2664.2005.01067.x | 2005 |
| *Lycaon pictus* | Canidae | Cross; Beissinger | Cons Biol | 10.1111/j.1523-1739.2001.00031.x | 2001 |
| *Lynx canadensis* | Felidae | Row; Wilson; Murray | J Anim Ecol | 10.1111/1365-2656.12179 | 2014 |
| *Lynx rufus* | Felidae | Schumaker; Ernst; White; Baker; Haggerty | Ecol Appl | 10.1890/02-5010 | 2004 |
| *Macaca mulatta* | Cercopithecidae | Hernandez-Pacheco; Rawlins; Kessler; Williams; Ruiz-Maldonado; Gonzalez-Martinez; Ruiz-Lambides; Sabat | Am J Primatol | 10.1002/ajp.22177 | 2013 |
| *Maccullochella peelii* | Percichthyidae | Yen; Bond; Shenton; Spring; Mac Nally | J Appl Ecol | 10.1111/1365-2664.12074 | 2013 |
| *Macquaria ambigua* | Percichthyidae | Yen; Bond; Shenton; Spring; Mac Nally | J Appl Ecol | 10.1111/1365-2664.12074 | 2013 |
| *Macrhybopsis storeriana* | Cyprinidae | Young; Koops | Fish & Oceans Can | NA | 2013c |
| *Macropus eugenii* | Macropodidae | Chambers; Bencini | Wildl Res | 10.1071/WR10080 | 2010 |
| *Malaclemys terrapin* | Emydidae | Mitro | Can J Zool | 10.1139/Z03-045 | 2003 |
| *Marmota flaviventris* | Sciuridae | Ozgul; Oli; Armitage; Blumstein; Van Vuren | Am Nat | 10.1086/597225 | 2009 |
| *Melanogrammus aeglefinus* | Gadidae | Gerber; Heppell | Biol Conserv | 10.1016/j.biocon.2004.01.029 | 2004 |
| *Membranipora membranacea* | Membraniporidae | Harvell; Caswell; Simpson | Oecologia | 10.1007/BF00323539 | 1990 |
| *Mesocentrotus franciscanus* | Strongylocentrotidae | Ebert; Russell | Mar Ecol Prog Ser | 10.3354/meps081031 | 1992 |
| *Micropterus dolomieu* | Centrarchidae | Spromberg; Birge | Environ Toxicol Chem | 10.1897/04-160.1 | 2005 |
| *Microtus oeconomus* | Muridae | Johannesen; Aars; Andreassen; Ims | Popul Ecol | 10.1007/s10144-003-0139-7 | 2003 |
| *Microtus sp.* | Muridae | Row; Wilson; Murray | J Anim Ecol | 10.1111/1365-2656.12179 | 2014 |
| *Milvus migrans* | Accipitridae | Sergio; Tavecchia; Blas; López; Tanferna; Hiraldo | Basic Appl Ecol | 10.1016/j.baae.2010.11.004 | 2011 |
| *Milvus milvus* | Accipitridae | Meyer; Meyer; Francisco; Holder; Verdonck | PLOS One | 10.1371/journal.pone.0147189 | 2016 |
| *Mirounga angustirostris* | Phocidae | Clinton; Le Boeuf | Ecology | 10.2307/1939945 | 1993 |
| *Mirounga leonina* | Phocidae | New; Clark; Costa; Fleishman; Hindell; Klanjsek; Lusseau; Kraus; McMahon; Robinson; Schick; Schwarz; Simmons; Thomas; Tyack; Harwood | Mar Ecol-Prog Ser | 10.3354/meps10547 | 2014 |
| *Montastraea annularis* | Faviidae | Hughes; Tanner | Ecology | 10.1890/0012-9658(2000)081[2250:RFLHAL]2.0.CO;2 | 2000 |
| *Morone saxatilis* | Moronidae | Gerber; Heppell | Biol Conserv | 10.1016/j.biocon.2004.01.029 | 2004 |
| *Moxostoma duquesnii* | Catostomidae | Velez-Espino; Koops | Ecol Model | 10.1016/j.ecolmodel.2012.09.022 | 2012 |
| *Moxostoma hubbsi* | Catostomidae | Velez-Espino; Koops | Ecol Model | 10.1016/j.ecolmodel.2012.09.022 | 2012 |
| *Mustela erminea* | Mustelidae | Wittmer; Powell; King | J Anim Ecol | 10.1111/j.1365-2656.2007.01274.x | 2007 |
| *Mya arenaria* | Myidae | Brousseau | Mar Biol | 10.1007/BF00390542 | 1978 |
| *Mytilus californianus* | Mytilidae | Carson; Cook; Lopez-Duarte; Levin | Ecology | 10.1890/11-0488.1 | 2011 |
| *Mytilus galloprovincialis* | Mytilidae | Carson; Cook; Lopez-Duarte; Levin | Ecology | 10.1890/11-0488.1 | 2011 |
| *Neogobius melanostomus* | Gobiidae | Spromberg; Birge | Environ Toxicol Chem | 10.1897/04-160.1 | 2005 |
| *Nephtys incisa* | Nephtyidae | Zajac; Whitlatch | Mar Ecol Prog Ser | 10.3354/meps057089 | 1989 |
| *Nocomis leptocephalus* | Cyprinidae | Peoples | Master Thesis | NA | 2010 |
| *Notropis anogenus* | Cyprinidae | Velez-Espino; Koops | Ecol Model | 10.1016/j.ecolmodel.2012.09.022 | 2012 |
| *Notropis percobromus* | Cyprinidae | Velez-Espino; Koops | Ecol Model | 10.1016/j.ecolmodel.2012.09.022 | 2012 |
| *Notropis photogenis* | Cyprinidae | Young; Koops | Fish & Oceans Can | NA | 2012a |
| *Noturus stigmosus* | Ictaluridae | Velez-Espino; Koops | Ecol Model | 10.1016/j.ecolmodel.2012.09.022 | 2012 |
| *Nuttallia obscurata* | Psammobiidae | Dudas; Dower; Anholt | Ecology | 10.1890/06-1216.1 | 2007 |
| *Odocoileus virginianus* | Cervidae | Edmunds | PhD Thesis | NA | 2013 |
| *Odocoileus virginianus subsp. borealis* | Cervidae | Jensen | Ecol Model | 10.1016/0304-3800(93)E0081-D | 1995 |
| *Oithona hebes* | Oithonidae | Torres-Sorando; Zacarias; Zoppi de Roa; Rodriguez | Ecol Model | 10.1016/S0304-3800(02)00355-1 | 2003 |
| *Oligocottus maculosus* | Cottidae | Pfister | Ecology | 10.1007/bf02027951 | 1996 |
| *Oncorhynchus clarkii* | Salmonidae | Velez-Espino; Koops | Ecol Model | 10.1016/j.ecolmodel.2012.09.022 | 2012 |
| *Oncorhynchus gilae* | Salmonidae | Brown; Echelle; Propst; Brooks; Fisher | Western N Am Naturalist | NA | 2001 |
| *Oncorhynchus kisutch* | Salmonidae | Spromberg; Birge | Environ Toxicol Chem | 10.1897/04-160.1 | 2005 |
| *Oncorhynchus tshawytscha* | Salmonidae | Wilson | Cons Biol | 10.1046/j.1523-1739.2003.01535.x | 2003 |
| *Onychogalea fraenata* | Macropodidae | Fisher; Hoyle; Blomberg | Ecol Appl | 10.2307/2641054 | 2000 |
| *Opsopoeodus emiliae* | Cyprinidae | Young; Koops | Fish & Oceans Can | NA | 2012b |
| *Orcinus orca* | Delphinidae | Vélez-Espino; Ford; Araujo; Ellis; Parken; Balcomb | Can Tech Report Fish & Aq Sci | 978-1-100-23563-9 | 2014 |
| *Oreamnos americanus* | Bovidae | Festa-Bianchet; Urquhart; Smith | Can J Zool | 10.1139/z94-004 | 1994 |
| *Osmerus spectrum* | Osmeridae | Velez-Espino; Koops | Ecol Model | 10.1016/j.ecolmodel.2012.09.022 | 2012 |
| *Ovis aries* | Bovidae | Clutton-Brock; Price; Albon; Jewell | J Anim Ecol | 10.2307/5330 | 1992 |
| *Ovis canadensis* | Bovidae | Rubin; Boyce; Caswell-Chen | J Wild Manag | 10.2307/3803144 | 2002 |
| *Ovis canadensis subsp. sierrae* | Bovidae | Johnson; Mills; Wehausen; Stephenson | Ecology | 10.1111/j.1365-2664.2010.01846.x | 2010 |
| *Pagurus longicarpus* | Paguridae | Damiani | Ecology | 10.1890/04-0956 | 2005 |
| *Palaemonetes pugio* | Palaemonidae | Manyin; Rowe | Aquat Toxicol | 10.1016/j.aquatox.2008.03.012 | 2008 |
| *Pan troglodytes subsp. schweinfurthii* | Hominidae | Morris; Altmann; Brockman; Cords; Fedigan; Pusey; Stoinski; Bronikowski; Alberts; Strier | Am Nat | 10.1086/657443 | 2011 |
| *Panopea generosa* | Hiatellidae | Ripley; Caswell | Popul Ecol | 10.1007/s10144-008-0075-7 | 2008 |
| *Panstrongylus geniculatus* | Reduviidae | Rabinovich; Feliciangeli | J Med Entomol | 10.1093/jme/tjv112 | 2015 |
| *Panthera pardus* | Felidae | Balme; Slotow; Hunter | Biol Cons | 10.1016/j.biocon.2009.06.020 | 2009 |
| *Papio cynocephalus* | Cercopithecidae | Morris; Altmann; Brockman; Cords; Fedigan; Pusey; Stoinski; Bronikowski; Alberts; Strier | Am Nat | 10.1086/657443 | 2011 |
| *Paramuricea clavata* | Plexauridae | Linares; Doak | Mar Ecol Pro Ser | 10.3354/meps08437 | 2010 |
| *Pelagia noctiluca* | Pelagiidae | Tomlinson; Maynou; Sabatés; Fuentes; Canepa; Sastre | Estuar Coast Shelf S | 10.1016/j.ecss.2015.11.012 | 2015 |
| *Percina copelandi* | Percidae | Velez-Espino; Koops | Ecol Model | 10.1016/j.ecolmodel.2012.09.022 | 2012 |
| *Perisoreus canadensis* | Corvidae | Schumaker; Ernst; White; Baker; Haggerty | Ecol Appl | 10.1890/02-5010 | 2004 |
| *Pernis apivorus* | Accipitridae | Bijlsma; Vermeulen; Hemerik; Klok | Ardea | 10.5253/078.100.0208 | 2012 |
| *Peromyscus maniculatus* | Muridae | Tallmon | PhD Thesis | NA | 2001 |
| *Petauroides volans* | Pseudocheiridae | Lindenmayer; Lacy; Pope | Ecol Appl | 10.2307/2641117 | 2000 |
| *Petaurus australis* | Petauridae | McCarthy; Lindenmayer; Possingham | Biol Cons | 10.1016/S0006-3207(00)00154-3 | 2001 |
| *Petromyzon marinus* | Petromyzontidae | Howe; Marsden; Donovan; Lamberson | J Great Lakes Res | 10.1016/j.jglr.2011.11.002 | 2012 |
| *Phacochoerus aethiopicus* | Suidae | Rodgers | Mammalia | 10.1515/mamm.1984.48.3.327 | 1984 |
| *Phalacrocorax auritus* | Phalacrocoracidae | Chastant; King; Weseloh; Moore | J Wild Manag | 10.1002/jwmg.628 | 2014 |
| *Phascolarctos cinereus* | Phascolarctidae | Baxter; McCarthy; Possingham; Menkhorst; McLean | Cons Biol | 10.1111/j.1523-1739.2006.00378.x | 2006 |
| *Phoca vitulina* | Phocidae | Heide-Jørgensen; Härkönen; Aberg | Ambio | 10.2307/4314005 | 1992 |
| *Phocarctos hookeri* | Otariidae | Lalas; Bradshaw | Biol Cons | 10.1016/S0006-3207(02)00421-4 | 2003 |
| *Phoebastria immutabilis* | Diomedeidae | Finkelstein; Doak; Nakagawa; Sievert; Klavitter | Anim Conserv | 10.1111/j.1469-1795.2009.00311.x | 2009 |
| *Phrynosoma cornutum* | Phrynosomatidae | Wolf; Hellgren; Schauber; Bogosian III; Kazmaier; Ruthven III; Moody | Popul Ecol | 10.1007/s10144-014-0450-5 | 2014 |
| *Physeter macrocephalus* | Physeteridae | Whitehead; Gero | Endangered Spp Res | 10.3354/esr00657 | 2015 |
| *Picoides arcticus* | Picidae | Rota; Millspaugh; Rumble; Lehman; Kesler | PLOS One | 10.1371/jourNAl.pone.0094700 | 2014 |
| *Picoides borealis* | Picidae | Maguire; Wilhere; Dong | J Wild Manag | 10.2307/3802460 | 1995 |
| *Pimephales promelas* | Cyprinidae | Gleason | Hum & Ecol Risk Assess | 10.1080/20018091094835 | 2001 |
| *Platygyra daedalea* | Faviidae | Riegl; Purkis | Glob Change Biol | 10.1111/gcb.13014 | 2015 |
| *Platygyra sinensis* | Faviidae | Babcock | Ecol Monog | 10.2307/2937107 | 1991 |
| *Plectus communis* | Plectidae | Kammenga; Van Gestel; Hornung | Ecol Appl | 10.2307/3061069 | 2001 |
| *Plexaura A* | Plexauridae | Lasker | Oecologia | 10.1007/BF00318316 | 1991 |
| *Pocillopora damicornis* | Pocilloporidae | Tanner | J Exp Mar Biol & Ecol | 10.1016/S0022-0981(97)00024-5 | 1997 |
| *Podocnemis expansa* | Podocnemididae | Mogollones; Rodríguez; Hernández; Barreto | Chel Cons Biol | 10.2744/CCB-0778.1 | 2010 |
| *Podocnemis lewyana* | Podocnemididae | Páez; Bock; Espinal-García; Rendón-Valencia; Alzate-Estrada; Cartagena-Otálvaro; Heppell | Copeia | 10.1643/CE-14-191 | 2015 |
| *Poecile atricapillus* | Paridae | Schumaker; Ernst; White; Baker; Haggerty | Ecol Appl | 10.1890/02-5010 | 2004 |
| *Poecilia reticulata* | Poeciliidae | Bronikowski; Clark; Rodd; Reznick | Ecology | 10.1890/0012-9658(2002)083[2194:PDCOPI]2.0.CO;2 | 2002 |
| *Polydesmus angustus* | Polydesmidae | David | Ann Entomol Soc Am | 10.1603/AN11151 | 2012 |
| *Pongo abelii* | Hominidae | Wich; Utami-Atmoko; Setia; Rijksen; Schürmann; van Hooff; van Schaik | J Hum Evol | 10.1016/j.jhevol.2004.08.006 | 2004 |
| *Porcellio scaber* | Porcellionidae | Kammenga; Van Gestel; Hornung | Ecol Appl | 10.2307/3061069 | 2001 |
| *Porites astreoides* | Poritidae | Edmunds | Mar Ecol Prog Ser | 10.3354/meps08595 | 2010 |
| *Porites harrisoni* | Poritidae | Riegl; Purkis | Glob Change Biol | 10.1111/gcb.13014 | 2015 |
| *Presbytis thomasi* | Cercopithecidae | Wich; Steenbeek; Sterck; Korstjens; Willems; Van Schaik | Am J Primatol | 10.1002/ajp.20386 | 2007 |
| *Proclossiana eunomia* | Nymphalidae | Radchuk; Turlure; Schtickzelle | J Anim Ecol | 10.1111/j.1365-2656.2012.02029.x | 2012 |
| *Procyon lotor* | Procyonidae | Schumaker; Ernst; White; Baker; Haggerty | Ecol Appl | 10.1890/02-5010 | 2004 |
| *Propithecus edwardsi* | Indriidae | Dunham; Erhart; Overdorff; Wright | 2008 | 10.1016/j.biocon.2007.10.006 | 2008 |
| *Propithecus verreauxi* | Indriidae | Morris; Altmann; Brockman; Cords; Fedigan; Pusey; Stoinski; Bronikowski; Alberts; Strier | Am Nat | 10.1086/657443 | 2011 |
| *Pseudocheirus peregrinus* | Pseudocheiridae | Lindenmayer; Lacy; Pope | Ecol Appl | 10.2307/2641117 | 2000 |
| *Pterois miles* | Scorpaenidae | Morris; Shertzer; Rice | Biol Invasions | 10.1007/s10530-010-9786-8 | 2011 |
| *Pterois volitans* | Scorpaenidae | Morris; Shertzer; Rice | Biol Invasions | 10.1007/s10530-010-9786-8 | 2011 |
| *Puffinus auricularis subsp. auricularis* | Procellariidae | Martinez-Gomez; Jacobsen | Biol Cons | 10.1016/S0006-3207(03)00171-X | 2004 |
| *Puffinus tenuirostris* | Procellariidae | Yearsley; Fletcher | Mat Biosci | 10.1016/S0025-5564(02)00119-0 | 2002 |
| *Puma concolor* | Felidae | Lambert; Wielgus; Robertson; Katnik; Cruickshank; Clarke; Almack | J Wildlife Manag | 10.2193/0022-541X(2006)70[246:CPDAVI]2.0.CO;2 | 2006 |
| *Pylodictis olivaris* | Ictaluridae | Sakaris; Irwin | Ecol Appl | 10.1890/08-0305.1 | 2010 |
| *Rana aurora* | Ranidae | Biek; Funk; Maxell; Mills | Conserv Biol | 10.1046/j.1523-1739.2002.00433.x | 2002 |
| *Rana catesbeiana* | Ranidae | Govindarajulu; Altwegg; Anholt | Ecol Appl | 10.1890/05-0486 | 2005 |
| *Rana temporaria* | Ranidae | Biek; Funk; Maxell; Mills | Conserv Biol | 10.1046/j.1523-1739.2002.00433.x | 2002 |
| *Rangifer tarandus* | Cervidae | Haskell; Ballard | J Wild Manag | 10.2193/2006-349 | 2007 |
| *Rangifer tarandus subsp. platyrhynchus* | Cervidae | Bjorkvoll; Lee; Grotan; Saether; Stien; Engen; Albon; Loe; Hansen | Ecol | 10.1890/15-0317.1 | 2016 |
| *Rangifer tarandus subsp. tarandus* | Cervidae | Messier; Huot; Le Henaff; Luttich | Arctic | 10.14430/arctic1733 | 1988 |
| *Rattus fuscipes* | Muridae | Lindenmayer; Lacy | Biol Cons | 10.1016/S0006-3207(01)00134-3 | 2002 |
| *Retropinna semoni* | Retropinnidae | Yen; Bond; Shenton; Spring; Mac Nally | J Appl Ecol | 10.1111/1365-2664.12074 | 2013 |
| *Rhinichthys cataractae* | Cyprinidae | Velez-Espino; Koops | Ecol Model | 10.1016/j.ecolmodel.2012.09.022 | 2012 |
| *Rhinichthys osculus* | Cyprinidae | Velez-Espino; Koops | Ecol Model | 10.1016/j.ecolmodel.2012.09.022 | 2012 |
| *Rutilus rutilus* | Cyprinidae | Otjacques; De Laender; Kestemont | Ecol Model | 10.1016/j.ecolmodel.2015.12.002 | 2016 |
| *Saguinus fuscicollis* | Callitrichidae | Watsa | PhD Thesis | 10.7936/K7DB7ZTD | 2013 |
| *Saguinus imperator* | Callitrichidae | Watsa | PhD Thesis | 10.7936/K7DB7ZTD | 2013 |
| *Salmo trutta* | Salmonidae | Fernández-Chacón; Genovart; Álvarez; Cano; Ojanguren; Rodriguez-Muñoz; Nicieza | Oecologia | 10.1007/s00442-015-3222-9 | 2015 |
| *Salvelinus confluentus* | Salmonidae | Bowerman | PhD Thesis | NA | 2013 |
| *Salvelinus fontinalis subsp. timagamiensis* | Salmonidae | Velez-Espino; Koops | Ecol Model | 10.1016/j.ecolmodel.2012.09.022 | 2012 |
| *Salvelinus malma* | Salmonidae | Spromberg; Birge | Environ Toxicol Chem | 10.1897/04-160.1 | 2005 |
| *Sardina pilchardus* | Clupeidae | Serghini; Boutayeb; Auger; Charouki; Ramzi; Ettahiri; Tchente | Acta Biotheor | 10.1007/s10441-009-9090-0 | 2009 |
| *Sceloporus arenicolus* | Phrynosomatidae | Ryberg; Hill; Painter; Fitzgerald | Cons Biol | 10.1111/cobi.12429 | 2014 |
| *Sceloporus grammicus* | Phrynosomatidae | Pérez-Mendoza | Herpetelogia | 10.1655/HERPETOLOGICA-D-12-00038R2 | 2013 |
| *Sceloporus mucronatus subsp. mucronatus* | Phrynosomatidae | Ortega-Leon; Smith; Zuniga-Vega; Mendez-de la Cruz | West N Am Naturalist | 10.3398/1527-0904(2007)67[492:GADOOP]2.0.CO;2 | 2007 |
| *Sceloporus woodi* | Lacertidae | Hokit; Stith; Branch | Cons Biol | 10.1046/j.1523-1739.2001.0150041102.x | 2002 |
| *Sciurus niger subsp. cinereus* | Sciuridae | Hilderbrand; Gardner; Ratnaswamy; Keller | Biol Cons | 10.1016/j.biocon.2007.01.015 | 2007 |
| *Scolytus ventralis* | Curculionidae | Berryman | Can Entomol | 10.4039/Ent1051465-11 | 1973 |
| *Setophaga cerulea* | Parulidae | Jones; Barg; Sillett; Veit; Robertson | Auk | 10.1642/0004-8038(2004)121[0015:MEOSAP]2.0.CO;2 | 2004 |
| *Sigmodon hispidus* | Muridae | Sauer; Slade | J Mamm | 10.2307/1381244 | 1985 |
| *Sparisoma viride* | Scaridae | O'Farrell; Salguero-Gómez; van Rooij; Mumby | J Anim Ecol | 10.1111/1365-2656.12399 | 2015 |
| *Spermophilus dauricus* | Sciuridae | Luo; Fox | J Mamm | 10.2307/1381947 | 1990 |
| *Spongia graminea* | Spongiidae | Cropper; Di Resta | Ecol Model | 10.1016/S0304-3800(99)00039-3 | 1999 |
| *Sprattus sprattus subsp. balticus* | Clupeidae | Haslob; Hauss; Petereit; Clemmesen; Kraus; Peck | Mar Biol | 10.1007/s00227-012-1933-6 | 2012 |
| *Stellifer illecebrosus* | Sciaenidae | Foster; Vincent | Aquat Conserv | 10.1002/aqc.2243 | 2012 |
| *Sterna hirundo* | Laridae | Szostek | J Orni | 10.1007/s10336-011-0745-7 | 2011 |
| *Sternotherus odoratus* | Kinosternidae | Mitchell | Herpetol Monogr | 10.2307/1467026 | 1988 |
| *Sternula antillarum subsp. browni* | Laridae | Massey; Bradley; Atwood | Condor | 10.2307/1369293 | 1992 |
| *Stratiodrilus aeglaphilus* | Histriobdellidae | Pardo; Vila; Bustamante | Hydrobio | 10.1007/s10750-007-9136-8 | 2008 |
| *Streblospio benedicti* | Spionidae | Levin; Caswell; DePatra; Creed | Ecology | 10.2307/1939879 | 1987 |
| *Strix occidentalis subsp. caurina* | Strigidae | Schumaker; Ernst; White; Baker; Haggerty | Ecol Appl | 10.1890/02-5010 | 2004 |
| *Strix occidentalis subsp. occidentalis* | Strigidae | LaHaye; Zimmermann; Gutiérrez | Auk | 10.1642/0004-8038(2004)121[1056:TVITVR]2.0.CO;2 | 2004 |
| *Sturnella neglecta* | Icteridae | Schumaker; Ernst; White; Baker; Haggerty | Ecol Appl | 10.1890/02-5010 | 2004 |
| *Sus scrofa subsp. scrofa* | Suidae | Gamelon; Besnard; Gaillard; Servanty; Baubet; Brandt; Gimenez | Evol | 10.1111/j.1558-5646.2011.01366.x | 2011 |
| *Tamiasciurus douglasii* | Sciuridae | Schumaker; Ernst; White; Baker; Haggerty | Ecol Appl | 10.1890/02-5010 | 2004 |
| *Tamiasciurus hudsonicus* | Sciuridae | McAdam; Boutin; Sykes; Humphries | EcoSci | 10.2980/1195-6860(2007)14[362:LHOFRS]2.0.CO;2 | 2007 |
| *Tautogolabrus adspersus* | Labridae | Gutjahr-Gobell; Zaroogian; Horowitz; Gleason; Mills | Ecotox & Env Sav | 10.1016/j.ecoenv.2005.05.017 | 2006 |
| *Thalassarche melanophris* | Diomedeidae | Arnold; Brault; Croxall | Ecol Appl | 10.1890/03-5340 | 2006 |
| *Thalia democratica* | Salpidae | Henschke; Smith; Everett; Suthers | J Planck Res | 10.1093/plankt/fbv024 | 2015 |
| *Trachemys scripta* | Emydidae | Frazer; Gibbons; Greene | Ecology | 10.2307/1447430 | 1990 |
| *Tribolium sp.* | Tenebrionidae | Constantino | Science | NA | 1997 |
| *Trichosurus caninus* | Phalangeridae | Lindenmayer; Lacy; Pope | Ecol Appl | 10.2307/2641117 | 2000 |
| *Tridacna gigas* | Cardiidae | Ripley; Caswell | Popul Ecol | 10.1007/s10144-008-0075-7 | 2008 |
| *Turdus torquatus* | Turdidae | Sim; Rebecca; Ludwig; Grant; Reid | J Anim Ecol | 10.1111/j.1365-2656.2010.01750.x | 2011 |
| *Tympanuchus cupido* | Phasianidae | Fefferman; Reed | J Wild Manag | 10.2307/3803419 | 2006 |
| *Umbonium costatum* | Trochidae | Noda; Nakao | J Anim Ecol | 10.2307/5722 | 1996 |
| *Upupa epops* | Upupidae | Gebreselassie | PhD Thesis | NA | 2010 |
| *Urocitellus armatus* | Sciuridae | Oli; Slade; Dobson | Ecology | 10.1890/0012-9658(2001)082[1921:EODROU]2.0.CO;2 | 2001 |
| *Urocitellus beldingi* | Sciuridae | Sherman; Morton | Ecology | 10.2307/1939140 | 1984 |
| *Urocitellus columbianus* | Sciuridae | Dobson; Oli | Am Nat | 10.1086/321322 | 2001 |
| *Urocyon littoralis* | Canidae | Hudgens; Garcelon | Oecologia | 10.1007/s00442-010-1761-7 | 2011 |
| *Ursus americanus* | Ursidae | Freedman; Portier; Sunquist | Ecol Model | 10.1016/S0304-3800(03)00171-6 | 2003 |
| *Ursus americanus subsp. floridanus* | Ursidae | Hostetler; McCown; Garrison; Neils; Barrett; Sunquist; Simek; Oli | Biol Cons | 10.1016/j.biocon.2009.05.029 | 2009 |
| *Ursus arctos* | Ursidae | Carroll; Noss; Paquet; Schumaker | Ecol Appl | 10.1890/02-5195 | 2003 |
| *Ursus arctos subsp. horribilis* | Ursidae | Pease; Mattson | Ecology | 10.2307/177030 | 1999 |
| *Ursus arctos subsp. yesoensis* | Ursidae | Kohira; Okada; Nakanishi; Yamanaka | Ursus | 10.2192/1537-6176-20.1.12 | 2009 |
| *Ursus maritimus* | Ursidae | Hunter; Caswell; Runge; Regehr; Amstrup; Stirling | Ecology | 10.1890/09-1641 | 2010 |
| *Vermivora chrysoptera* | Parulidae | Bulluck; Buehler; Vallender; Robertson | Wilson J Ornithol | 10.1676/12-154.1 | 2013 |
| *Vipera aspis* | Viperidae | Altwegg; Dummermuth; Anholt; Flatt | Oikos | 10.1111/j.0030-1299.2001.13723.x | 2005 |
| *Vireo latimeri* | Vireonidae | Woodworth | Cons Biol | 10.1046/j.1523-1739.1999.97267.x | 1999 |
| *Vulpes vulpes* | Canidae | McLeod; Saunders | Wild Res | 10.1071/WR00104 | 2001 |
| *Watersipora subtorquata* | Watersiporidae | Cockrell; Sorte | J Exp Mar Bio & Ecol | 10.1016/j.jembe.2012.11.010 | 2013 |
| *Xenosaurus grandis* | Xenosauridae | Zuniga-Vega; Valverde; Rojas-Gonzalez; Lemos-Espinal | Copeia | 10.1643/0045-8511(2007)7[324:AOTPDO]2.0.CO;2 | 2007 |
| *Xenosaurus platyceps* | Xenosauridae | Rojas-Gonzalez; Jones; Zúñiga-Vega; Lemos-Espinal | Amphibia-Reptilia | 10.1163/156853808784124992 | 2008 |
| *Xenosaurus sp.* | Xenosauridae | Zamora-Abrego; Chang; Zuniga-Vega; Nieto-Montes de Oca; Johnson | Herpet | 10.1655/09-005.1 | 2010 |
| *Xestospongia muta* | Petrosiidae | McMurray; Henkel; Pawlik | Ecology | 10.3354/meps339093 | 2010 |
| *Yoldia notabilis* | Yoldiidae | Nakaoka | Oikos | 10.2307/3546090 | 1997 |
| *Zalophus californianus* | Otariidae | Wielgus; Gonzalez-Suarez; Aurioles-Gamboa; Gerber | Ecol Appl | 10.1890/07-0892.1 | 2008 |
| *Zenaida macroura* | Columbidae | Schumaker; Ernst; White; Baker; Haggerty | Ecol Appl | 10.1890/02-5010 | 2004 |
| *Zingel asper* | Percidae | Labonne; Gaudin | Can J Fish Aq Sci | 10.1139/f05-245 | 2006 |
| *Ziphiidae sp.* | Ziphiidae | Chiquet; Montgomery; Ma; Ackleh | Neu Par Sci Comp | 10.1002/jwmg.835 | 2015 |
| *Zoarces viviparus* | Zoarcidae | Bergek; Ma; Vetemaa; Franzén; Appelberg | Ecotoxicol Environ Safety | 10.1016/j.ecoenv.2012.01.019 | 2012 |
| *Zonotrichia leucophrys* | Emberizidae | Baker; Mewaldt; Stewart | Ecology | 10.2307/1937731 | 1981 |
| *Zootoca vivipara* | Lacertidae | Mugabo; Perret; Legendre; Le Galliard | J Anim Ecol | 10.1111/1365-2656.12109 | 2013 |

**Table S2.** Plant traits linked to modularity used in our meta-analysis. We reviewed nine plant traits (five aboveground, four belowground traits) in plants for overlap with demographic measures in the COMADRE database in order to identify whether these traits relate to senescence in reflection of Finch’s Hypothesis that modularity influences senescence. In total, 138 plants were represented across datasets.

| **Traits linked to modularity** | **Database** | **Number of overlapping species with COMPADRE** |
| --- | --- | --- |
| Leaf life span | BIEN | 43 |
| Maximum whole plant longevity | BIEN | 109 |
| Stem wood density | BIEN | 98 |
| Vessel lumen area | BIEN | 30 |
| Vessel number | BIEN | 30 |
| Belowground biomass per ground area | FRED | 25 |
| Belowground biomass per plant | FRED | 23 |
| Root diameter | FRED | 42 |
| Root lignin content | FRED | 28 |

**Table S3**. List of traits linked to modularity in animals. This table outlines traits of each major physiological system in animals relevant to modularity and whether they offer supporting evidence or counterevidence for modularized function. The traits and summary determinations described were based on a comprehensive review of comparative physiological texts and review papers and provided the high-level framework for triaging what systems were plausibly modularized in function, and meriting of further attention within the scope of our study. The immune and renal systems were selected for further study in our meta-analysis.

| **System** | **Likelihood of Modularity** | **Supporting evidence** |
| --- | --- | --- |
| Lymphatic System | Plausible | [*For*] Lymph nodes (spatially discrete, functionally identical); multiple, inter-dependent functional components: spleen (numbering up to 14 in cetaceans; multiple constituents in sharks), thymus, and bone marrow.  [*Against*] Continuity of lymph circulation. |
| Immune System | Plausible | [*For*] Multiple organs active in the role of ionic exchange (gills, kidneys, salt-glands); partial independence of lobes of a reniculated kidney; multiple kidneys in most mammals.  [*Against*] Little variation in the number of kidneys or other osmoregulatory organs. |
| Cardiovascular System | Plausible | [*For*] Vessel multiplicity; potential for multiple vascular pathways to provide circulation to the same area (i.e. collateral vasculature).  [*Against*] Vasculature is highly integrated and structurally contiguous (cf. spatially discrete repeat units). |
| Skeletal System | Implausible | [*For*] Highly discrete structures.  [Against] Strong interdependency. Few cases of structural redundancy have been identified in the literature. |
| Neurological Systems | Implausible | [*Against*] Enervation seldom involves multiple neurons that retain full functional integrity if another is removed or functionally immobilized. |
| Endocrine systems | Implausible | [*For*] Routine functional overlap of efferent signals.  [*Against*] Multiple physiological responses associated with most hormones complicates the specificity on which redundancy can be understood under a traditional sectorial framework. |
| Respiratory | Implausible | [*For*] Discrete, repeat units of respiratory pigments.  [*Against*] Known respiratory surfaces do not have discrete tissue divisions which structurally or functionally sub-divide the organ(s) of interest. No known evidence of how injury to one population of respiratory pigments could be off-set by another. |
| Dermal | Implausible | [*Against*] Similar to respiratory exchange organs, there are not clear boundaries in the tissue that would potentially serve to structurally or functionally subdivide the intermediating organs of interest. |
